# An Adaptive H-Refinement Method for the Boundary Element Fast Multipole Method for Quasi-static Electromagnetic Modeling

**DOI:** 10.1101/2023.08.11.552996

**Authors:** William A Wartman, Konstantin Weise, Manas Rachh, Leah Morales, Zhi-De Deng, Aapo Nummenmaa, Sergey N Makaroff

## Abstract

**Objective:** In our recent work pertinent to modeling of brain stimulation and neurophysiological recordings, substantial modeling errors in the computed electric field and potential have sometimes been observed for standard multi-compartment head models. The goal of this study is to quantify those errors and, further, eliminate them through an adaptive mesh refinement (AMR) algorithm. The study concentrates on transcranial magnetic stimulation (TMS), transcranial electrical stimulation (TES), and electroencephalography (EEG) forward problems.

**Approach:** We propose, describe, and systematically investigate an AMR method using the Boundary Element Method with Fast Multipole Acceleration (BEM-FMM) as the base numerical solver. The goal is to efficiently allocate additional unknowns to critical areas of the model, where they will best improve solution accuracy.

The implemented AMR method’s accuracy improvement is measured on head models constructed from 16 Human Connectome Project subjects under problem classes of TES, TMS, and EEG. Errors are computed between three solutions: an initial non-adaptive solution, a solution found after applying AMR with a conservative refinement rate, and a “silver-standard” solution found by subsequent 4:1 global refinement of the adaptively-refined model.

**Main Results:** Excellent agreement is shown between the adaptively-refined and silver-standard solutions for standard head models. AMR is found to be vital for accurate modeling of TES and EEG forward problems for standard models: an increase of less than 25% (on average) in number of mesh elements for these problems, efficiently allocated by AMR, exposes electric field/potential errors exceeding 60% (on average) in the solution for the unrefined models.

**Significance:** This error has especially important implications for TES dosing prediction – where the stimula t ion strength plays a central role – and for EEG lead fields. Though the specific form of the AMR method described here is implemented for the BEM-FMM, we expect that AMR is applicable and even required for accurate electromagnetic simulations by other numerical modeling packages as well.

## 1. Introduction

In the last decade, noninvasive electrical neurostimulation methods have been increasingly popular topics of research and application for a wide variety of psychiatric disorders. Transcranial magnetic stimulation (TMS), in which an electromagnetic coil placed on the scalp induces electrical currents in the brain, has been applied to depression [1], anxiety [2], and addiction [3], among other uses. Transcranial electrical stimulation (TES), in which electrodes placed on the scalp inject current directly through the intervening tissues into the brain, has been applied to study problems including Alzheimer’s Disease [4], depression [5], and epilepsy [6]. Electroencephalography (EEG) has been applied in conjunction with both TMS and TES [4][5][6][7] to quantify and localize neuronal responses to stimulation by these methods.

As these methods have been applied to more problems and with greater requirements for precision, pre-stimulation planning and post-stimulation analysis have become vital components of experimental design. The planning and analysis both have been relying increasingly on accurate numerical electromagnetic analysis by open-source packages such as SimNIBS [8], ROAST [9], and BEM-FMM [10]. Such methods, however, are only as accurate as the computational models upon which they operate – frequently segmented from MRI data by medical image processing packages such as *headreco* [30]. In 2020, Gomez et. al. [16] carried out an investigation on, among other parameters, the necessary computational mesh resolution to accurately simulate TMS trials under various electromagnetic solver formulations. The process was time and attention intensive, requiring construction of progressively higher and higher resolution meshes and comparison against a presumed-accurate result. The meshes in this study were refined globally, resulting in a 4x increase in number of surface elements per refinement level or a staggering 8x increase in number of volumetric elements per refinement level. These are steep prices to pay for guarantees of accurate simulation, but are the only available means to achieve such a guarantee without a more efficient mechanism.

To this end, we introduce, investigate, and describe a fast, automated adaptive mesh refinement (AMR) method applicable to TES, TMS, and EEG modeling problems. AMR is understood as an automated local refinement of a computational mesh in domains where the discretization error is highest. It is repeated until a user-specified convergence criterion (e.g. relative error between two iterations becomes less than 0.1%) or termination criterion (30 AMR steps elapsed) is met. AMR is a chief feature of high-end commercial ANSYS FEM (Finite Element Method) software for demanding low-frequency, high-frequency, and power applications [15]. The method is implemented and tested as an extension of our Boundary Element Fast Multipole Method (BEM-FMM), which has been applied to model all three mentioned stimulation/recording modalities [10][11][12] in addition to other problems [13][14]. To our knowledge, none of the other major electromagnetic modeling packages for electromagnetic brain stimulation offer an AMR method to date.

Refinement can take two main forms: a geometric bisection of a given element (“h-refinement”) or an increase of the local approximation order (“p-refinement”). In this study, we consider only h-refinement. Given an initial finite element approximation, the basic idea of an h-adaptive method is to create a refined partition by subdividing those elements where local error estimators indicate that the error is large; the next approximation to the solution is computed using the newly created model, and the process repeats. Because of their success in practice, the use of such adaptive methods has become more widely spread in recent years [17][18][19][20]. For the boundary element method (BEM), a similar methodology applies [21][22].

In contrast to FEM-based solvers, BEM-FMM is capable of unconstrained numer ica l resolution. It is capable of computing the electric field and potential (or pseudo-potential) at any given observation points, including points not known a priori and points arbitrarily close to model interfaces [10]. Where FEM-based methods must introduce additional volumetric elements to support such observation points in regions of rapidly-varying E-field, the BEM-FMM can compute a non-interpolated result that is nearly exact for the given (inexact) model geometry and the zeroth-order charge density residing upon it. Adaptive mesh refinement for BEM-FMM was initia l ly introduced in [27] to accurately determine the effects of thin meningeal layers on TMS and TES problems. However, no systematic investigation of the method had yet been carried out and no other applications except for meningeal layers have been considered.

In this work, we introduce and describe the full implementation of an efficient adaptive mesh refinement algorithm for BEM-FMM. We systematically evaluate the accuracy improvement achieved due to AMR for TMS, TES, and EEG forward problem classes on realistic human head models. The source code is available for download in an OSF repository [23].

## 2. Materials and Methods

### 2.1. Charge-Based Formulation of the Boundary Element Fast Multipole Method

For TMS, TES following a current-based electrode approximation, and EEG, the BEM-FMM is formulated as a Fredholm equation of the second kind [10][12]. For TES problems following a voltage-based electrode approximation, the method additionally incorporates a Fredholm equation of the first kind via weak (additive) coupling [11].

The present BEM-FMM makes several assumptions about the problem being modeled. First, it assumes that the model can be divided into compartments of homogeneous, linear, isotropic, conductive media. Second, it assumes that any electromagnetic waves have a very long wavelength compared to the model dimensions, so that the problem is quasi-static in nature. Third, it assumes that any secondary magnetic fields are negligible in magnitude compared to any primary magnetic fields.

Following the assumption that compartments of the model are conductive, electric charges cannot accumulate in the volume. They must instead accumulate as surface charges at interfaces (boundaries, surfaces) between materials of different conductivities. Under the quasi-static assumption, these accumulated surface charges are sufficient to fully characterize (via Coulomb’s Law) the secondary electric field ***E***^*S*^(***r***), which can be added to the primary electric field ***E***^*P*^(***r***) to recover the total electric field ***E***(***r***) = ***E***^*P*^(***r***) + ***E***^*S*^(***r***) at any arbitrary observation point ***r*** inside, outside, or on a surface of the model. The electric potential *V*(***r***) can be similarly recovered at any arbitrary observation point.

The BEM-FMM solution procedure is carried out in two main steps to most efficiently utilize the FMM. The first step is to solve for the charge density *ρ(****r***) that arises on interfaces between different materials due to a primary (external) electric field or enforced electric potential. The second step is to recover field quantities of interest (e.g. electric field, voltage, current density) at any observation points in terms of the primary field and the surface charges induced by that primary field.

Eq. (1) (cf. [10][1]) is the continuous form of the integral equation when the excitation can be written as a primary (external) electric field ***E***^*p*^(***r***).

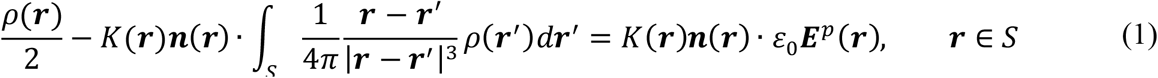

Here, *S* is the set of all points (ℝ^3^) lying on any boundary (surface) *S* between two materials of different properties, ***r*** is an arbitrary point on a boundary, ρ(***r***) is the surface charge density at ***r***, 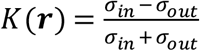 is the contrast between the conductivity just inside (σ_*in*_) and just outside (σ_*out*_) the boundary on which ***r*** lies, ***n***(***r***) is the unit vector normal to the boundary at ***r***, ε_0_ is the permittivity of free space, and ***E***^*p*^(***r***) is the primary electric field incident on the boundary at ***r***. This equation is to be solved for the surface charge density ρ(***r***).

For TMS, the primary electric field ***E***^*p*^(***r***) on the right-hand side of Eq. (1) can be written in terms of the magnetic vector potential applied by the coil when driven by a time-varying electric current as described in Appendix A of [10]. For TES electrodes that are assumed to inject a uniform current flux density over their area, the primary electric field can be written in terms of the injected current density and the conductivity of the interior tissue (and set to zero for any facet that does not touch an electrode) [11]. For EEG, the primary field radiates from clusters of charge dipoles and can be evaluated by FMM-accelerated application of Coulomb’s Law [12].

Certain problems modeled by the BEM-FMM cannot be straightforwardly written in terms of a primary electric field. For example, if electrodes in a TES problem are assumed to maintain constant electric potentials on their surfaces, then the injected current flux density is not necessarily spatially constant and cannot be used to estimate a primary electric field. In this case, the primary electric field is 0 everywhere, and an additional constraint (specified in Eq. (2) below) is additively coupled into the integral system:

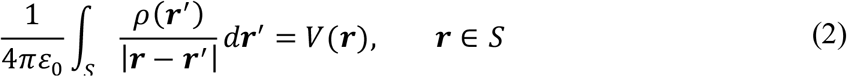

Here, *V*(***r***) denotes the electric potential that is externally enforced at ***r*** on any boundary. For locations that do not touch electrodes or otherwise have an enforced voltage, *V*(***r***) = 0. This is a Fredholm equation of the first kind.

The head model is constructed as a collection of triangular surface meshes representing the boundaries between different tissues. The charge density is expanded in terms of zeroth-order (pulse) basis functions – in other words, the charge density *c*_*m*_ is assumed to be constant over the entire surface of any individual facet *m*, but may vary facet-to-facet. Discretizing Eq. (1) via the Galerkin method, Eq. (3) is obtained.

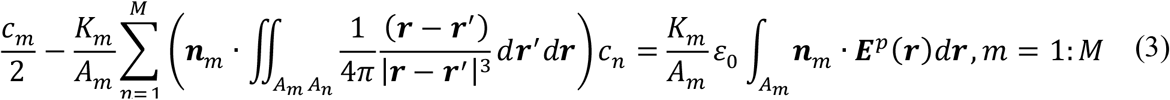

In Eq. (3), *M* denotes the total number of triangular surface elements in the model. Eq. (3) can be rewritten in matrix form as

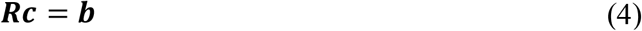

Note that the matrix ***R*** is never explicitly constructed in the BEM-FMM; instead, the FMM is applied in conjunction with a sparse near-field correction to directly compute the matrix-vector product ***Rc*** when necessary. The on-diagonal elements 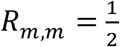 describe the self-interaction of the planar charge density on triangle *m*. The off-diagonal elements ***R***_*m*,*n*_ describe the average normal component of the E-field contributed to triangle *m* by a charge density *c*_*n*_ residing on triangle *n*:

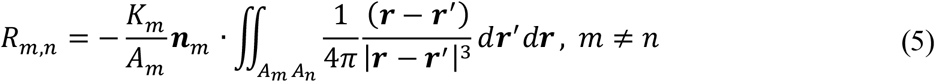

For triangles sufficiently distant (> 2 to 5 average triangle radii) from each other, ***R***_*m*,*n*_ can be approximated as:

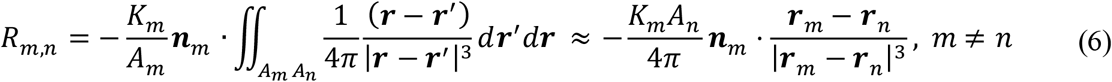

where ***r***_*m*_ and ***r***_*n*_ denote the respective centroids of triangles *m* and *n*. Interactions of this form can be accelerated dramatically by the FMM. For triangles close to each other, the full double integral over both triangles must be precomputed and applied as a correction to the FMM-accelerated initial computation.

In the matrix equation formulation, the elements *b*_*m*_ of ***b*** are straightforwardly given by

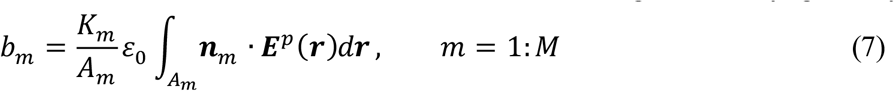

The system is solved iteratively for ***c*** using the Generalized Minimum Residual Method (GMRES). Once the charge solution ***c*** is known, the electric field can be recovered at any observation point not residing directly on a model surface according to Coulomb’s law Eq. (8):

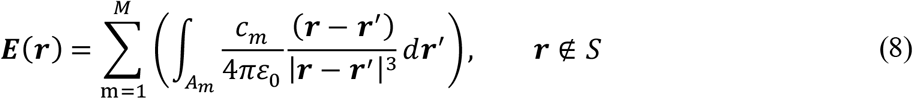

This computation can be similarly accelerated via the FMM.

### 2.2 Accuracy limit: 0^th^ order (pulse) basis functions

As stated, the BEM-FMM solves for the charge density that accumulates at interfaces between tissues of differing conductivities. From this charge density, any quantities of interest (e.g. electric field, current density, or electric potential) can be recovered at arbitrarily-positioned observation points, including observation points very close to or lying on the charged interfaces. Under the assumption that the BEM-FMM has produced a physically realistic and accurate charge distribution, the desired quantities can be computed at arbitrary observation points with high accuracy.

The assumption of a realistic charge distribution may sometimes be violated due to the BEM-FMM’s use of zeroth order (pulse) basis functions. These basis functions effectively hold the charge density spatially constant over the area of any given triangle. In complex regions of the model – for example, in regions of sharp curvature or with multiple boundaries in close proximity – the initial mesh may not provide enough facets to support a charge density that varies sufficiently rapidly in a spatial sense. Such infidelity can give rise to multiple sources of error. The more benign is a local error that affects accuracy of field calculations at observation points in the vicinity of the complex region. The more insidious is error that affects triangle-to-triangle interactions during the solver’s iterative phase, as this error can propagate to distant regions of the model. As will be shown, EEG and TES forward problems are particularly susceptible to this latter error because their primary electric fields are localized to small regions and depend on triangle-to-triangle interactions to propagate their effects through the model.

### 2.3 Algorithmic Description: Adaptive Mesh Refinement Applied to BEM-FMM

To mitigate the aforementioned shortcoming of the zeroth order basis functions, an AMR scheme was incorporated into the BEM-FMM. This scheme is based on *h*-refinement, meaning that it operates by subdividing existing mesh elements into a larger number of smaller elements. It aims to improve the quality of the initial charge solution and subsequent electric field reconstructions by selectively increasing the mesh resolution in critical or complex areas of the model, without unnecessarily increasing mesh resolution in areas experiencing fields or charge densities with low spatial variation. It efficiently allocates additional degrees of freedom in the locations where they will best improve solution accuracy.

As currently implemented, the BEM-FMM augmented by AMR is carried out in alternating steps of “Solve” and “Refine”. During the “Solve” step, the incident stimulus/constraints are applied to the current version of the model, and the charge solution is obtained using an iterative solver (GMRES). To preserve existing solution progress, the final charge density solution ***c*** from the prior model step is chosen as the initial estimate for the charge density solution for the current model. During the “Refine” step, the current model and solution are evaluated, certain facets are selected for subdivision, and neighbor integrals are recomputed. The “Refine” step creates a new model that has the same geometry as the prior model but introduces a greater number of unknowns.

Facets are selected for subdivision according to the *total* charge upon them (i.e., *Q*_*m*_ = *c*_*m*_*A*_*m*_, where *A*_*m*_ is the area of facet *m*). For each surface in the model, a user-specified proportion ***r*** of facets belonging to that surface are selected for refinement in order of highest to lowest absolute value of total charge. This allocation on a per-surface basis distributes the locations of refinement throughout the model, as otherwise they would tend to be allocated exclusively to the location of the strongest source (e.g. TES electrodes). Distributing the locations of refinement in this manner helps smooth the convergence of the solution and prevent instances where the error function reaches a local minimum.

Refinement is performed via simple barycentric subdivision, wherein three additional vertices are inserted at the midpoints of each selected triangle’s edges to break it into four sub-triangles. These new triangles are coplanar with and similar to the original triangle. No remeshing operation needs to be performed to restore mesh connectivity or manifoldness, as the BEM-FMM with zeroth-order pulse bases is unaffected by mesh manifoldness or lack thereof. This is a chief advantage of the proposed method.

The new triangles inherit the charge density of the original triangle to preserve solution progress. The charge density is not scaled upon inheritance since it is by definition already expressed per unit area. Fig. 1 shows an example of a small region of a mesh after multiple adaptive refinement steps.

**Fig. 1:**
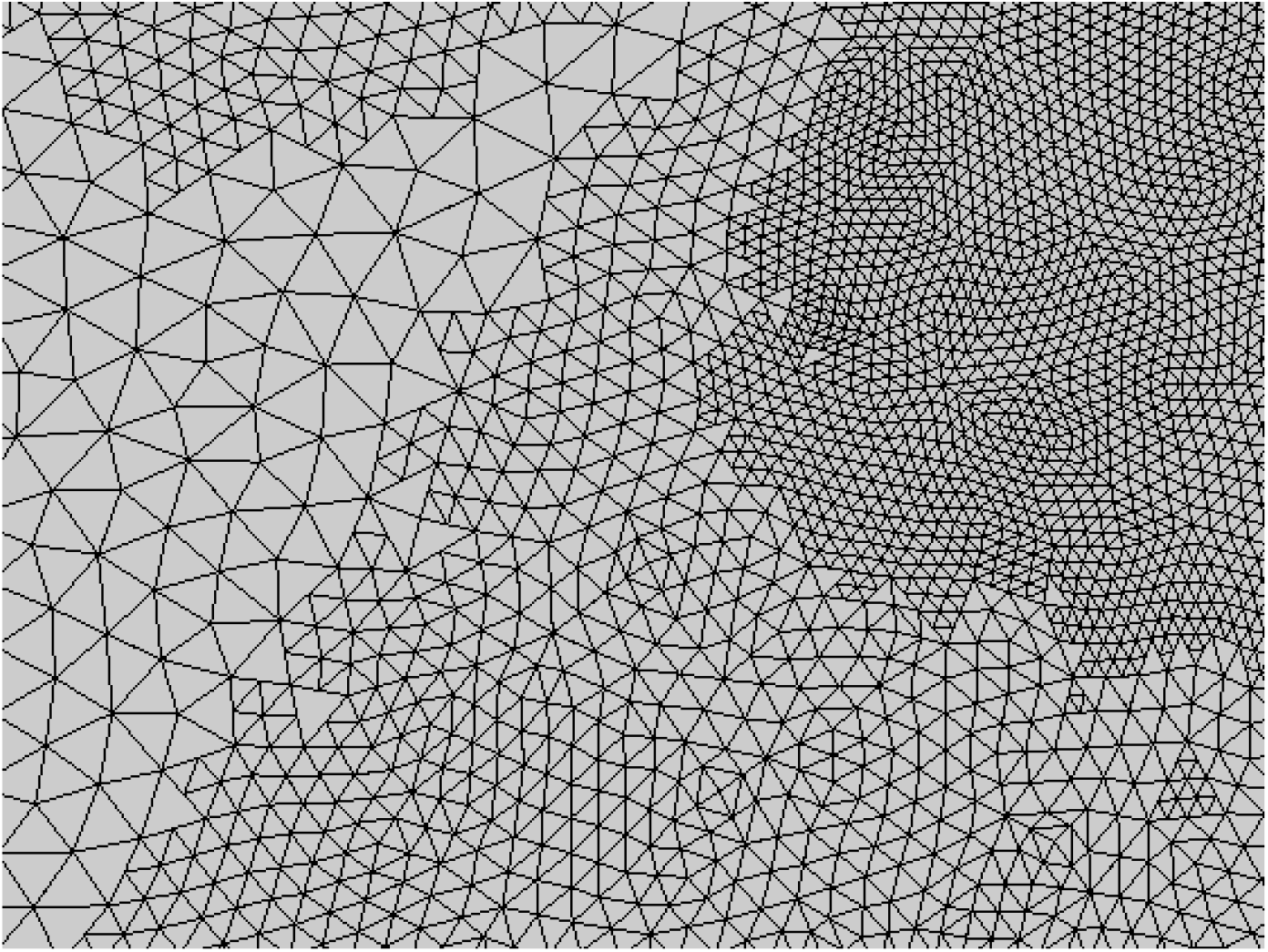
Example of a small region of mesh subjected to adaptive mesh refinement in the vicinity of a focal current source (out of frame past the upper right corner). Three distinct levels of subdivision are visible: unmodified facets of the initial mesh (left), facets subjected to one AMR pass (center, top left, and bottom right), and facets subjected to two AMR passes (top right). No attempt is made to restore mesh connectivity across neighboring triangles subjected to different levels of refinement since no BEM-FMM basis functions depend on such connectivity.

Multiple termination criteria can be defined for the adaptive refinement method. A natural termination condition may involve, for example, monitoring the convergence of the electric field in a predefined region of interest (ROI) and terminating when the relative error between adaptive steps drops below a predefined threshold. This is the termination condition used in this study. Another natural metric is the error between the charge density (solution) vectors produced by subsequent AMR steps, which would decrease the computational time required by removing the need to carry out (e.g.) an E-field recovery step after every AMR pass.

### 2.4 Expected Performance of Adaptive Mesh Refinement

If ***r*** denotes the refinement rate (fraction of facets refined per adaptive pass) and *k* denotes the number of adaptive passes applied, then the number of facets (unknowns) in the mesh after refinement is given by:

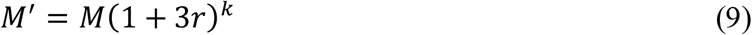

Where *M* and *M*′ denote the total number of facets pre- and post-refinement, respectively. If the average mesh edge length in the model pre-refinement is denoted by *l*, then the edge length *l*′ of an average facet subjected to maximum possible refinement is given simply by

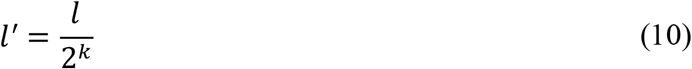

Eqs. 9 and 10 show that this implementation of adaptive mesh refinement grows the model exponentially and is capable of increasing the mesh resolution exponentially in critical regions. If the number of AMR iterations were to approach infinity, it is expected that all discrete charges *Q*_*m*_ present in the model would be drawn equal to each other.

To put these equations into perspective with reasonable values of *k* and ***r*** (used during preliminary investigations for TES and EEG), consider Connectome Subject 110411 of the Human Connectome Project [28][29] meshed by the commonly used *headreco* pipeline [30]. Pre-refinement, this model has 1.04 M facets and average edge length 1.44 mm. After 16 adaptive refinement steps at a refinement rate of 1% per step (*k* = 16; ***r*** = 0.01), the number of facets (unknowns) would increase by 60.5% to 1.67 M facets. If a certain average facet in a critical region were subdivided on every adaptive pass, its edge length would be scaled by a factor of 1.526e-5, resulting in a final edge length of 22.0 nm.

By contrast, suppose one iteration of global barycentric subdivision were applied. The mesh size would increase by 300% (total 4.16 M facets), but the edge length would only be scaled by a factor of 0.5: an average edge would decrease from 1.44 mm to 0.72 mm. Compared to global refinement, adaptive mesh refinement applied in this format is capable of increasing the mesh resolution to very high levels in vital regions while allocating the new unknowns efficiently.

### 2.5 Human Head Models Under Test

The human head models considered in the following accuracy tests are 16 subjects from the Human Connectome Project [28][29]. Surface mesh models for these subjects were generated using the *headreco* pipeline [30]. These models have 1.06 M facets on average, have an average triangle edge length of 1.43 mm, and contain seven tissues. The tissues are air, skin, skull, cerebrospinal fluid, gray matter, white matter, ventricles, and eyes as shown in Fig. 2 c,d. In general, we refer to a given boundary by the name of its interior tissue. For example, the “gray matter surface” would refer to the GM/CSF boundary.

**Fig. 2:**
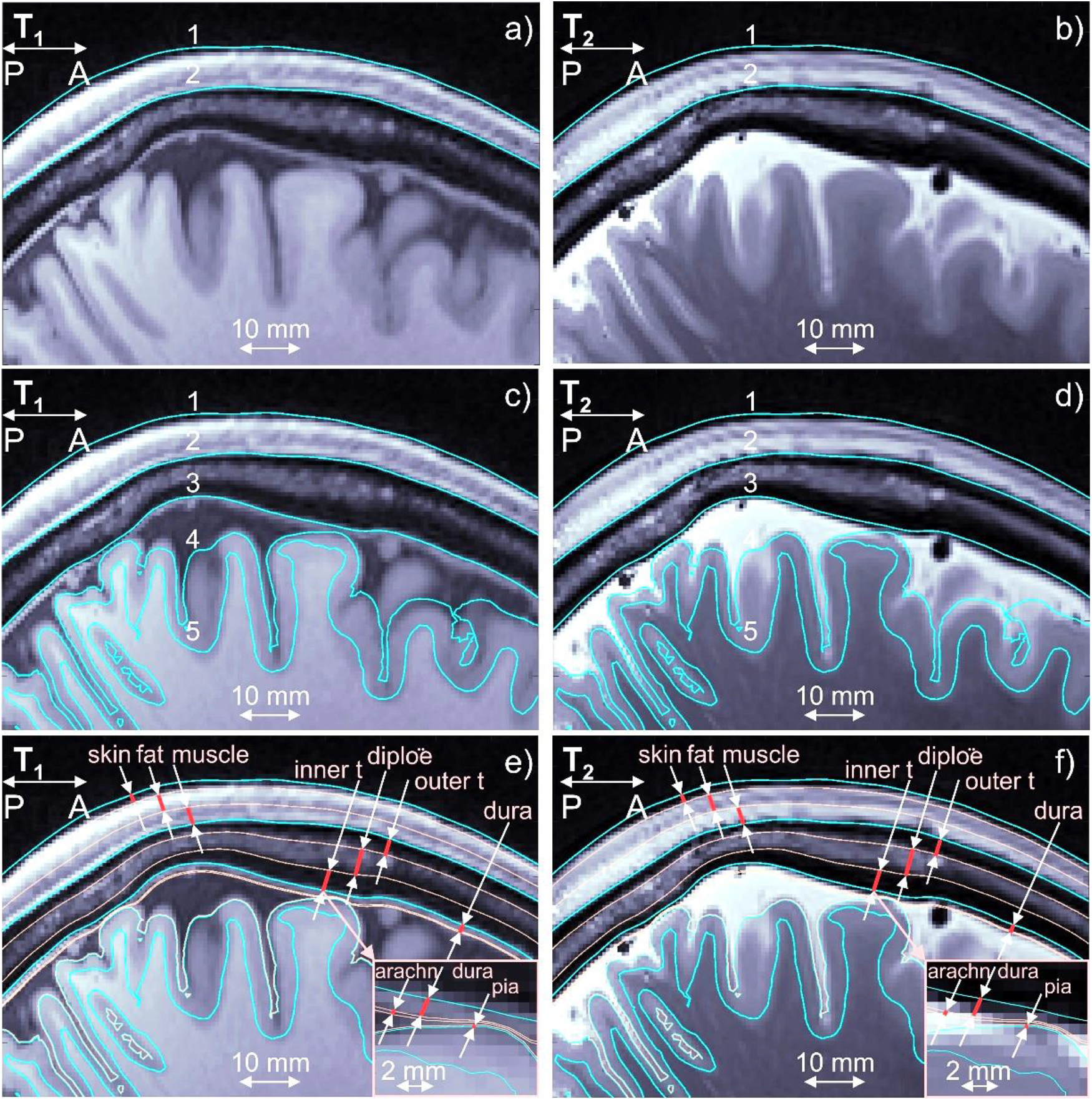
a,b) T1/T2 Images for Connectome subject 120111 and headreco segmentation for scalp (1) and skull (2) shown in blue. Dura mater is seen on the T1 image. c,d) The same images and base headreco segmentation for scalp (1), skull (2), CSF (3), gray matter (4), and white matter (5). The headreco routine subsumes the dura mater into the CSF volume. e,f) Base headreco segmentation (blue) and new extracerebral compartments (pale pink). They agree with the background MRI information. Two insets display meninges.

A second set of models was derived based on these 16 Connectome subject models in the spirit of our prior work [27], where we investigated the impact of meningeal layers that are not commonly segmented by the major packages. These 16 extended models contain additional tissue boundaries constructed by expansion or contraction of existing boundaries as shown in Fig. 2e,f. The skin volume was subdivided into layers of skin, fat, and muscle. The skull volume was subdivided into two layers of cortical bone separated by one layer of cancellous (spongy) bone. Layers representing the dura mater, arachnoid mater, and pia mater were introduced in the CSF volume outside the GM. In this study, we are proposing to *add to existing segmentations* skin, fat, and muscle of the scalp, outer table, diploë, and inner table of the skull, and three brain meninges, all via known anatomical rules:

#### Scalp→skin, fat, muscle

To partition space between skin and bone shells, the following data can be used: skin – 20%, fat – 40%, muscle – 40%. These values are widely used in safety studies for MR RF coils ([34] and other sources cited there). Other references (e.g., [33]) predict the conductivities.

#### Skull→outer table, diploë, inner table

Based on data from [35]-[39] for 300 subjects, the following estimates can be deduced in the frontal lobe: outer table (cortical) – 30%; diploë (cancellous) – 40%; inner table (cortical) – 30%. For the parietal lobe, the diploë thickness may exceed 50% [39]. The following values can be used there: outer table – 30%; diploë – 50%; inner table – 20%. A smooth transition is automatically made from one lobe to another. Variations of this scheme are easily programmable. The conductivity values from [31] can be used: cortical bone: 6.4 mS/m, cancellous bone: 29 mS/m.

#### CSF→dura mater, arachnoid, true CSF, pia mater

Here, we can use integral data given in [32], [40]-[43]: 1.11 mm for dura, 0.2 mm for arachnoid, and 0.1 mm for pia mater except in the longitudinal fissure. Further algorithmic details are given in [27]. The final models have 1.59 M facets on average with an average triangle edge length of 1.45 mm.

It is critical to note that these 14-tissue models are introduced for the sole purpose of testing the AMR method, and not for the purpose of comparing their solutions against their corresponding 7-tissue models. Based on previous work [27], it is expected that the extra tissues have a substantial impact on TES and EEG simulations, and that accurate segmentation of these tissues will be necessary for future applications. The construction of these tissues based on anatomical rules represents an attempt to characterize the solvability and convergence of this future class of problem. The solutions themselves may be inaccurate due to the conjectural nature of the models.

Tables 1 and 2 contain the conductivities assigned to each tissue type appearing in the 14-tissue and 7-tissue models, respectively. Entries marked by an asterisk (*) denote values that were computed by a weighted average of composite tissues’ individual conductivities. The “Skin” conductivity for the 7-tissue model was computed by a weighted average of the “Skin”, “Fat”, and “Muscle” conductivities from the 14-tissue model with weights assigned according to relative thicknesses of these layers. Similarly, the “Bone” conductivity for the 7-tissue model was computed by a weighted average of the “Cortical Bone” and “Trabecular Bone” tissues.

**Table 1:** Tissue Conductivities Assigned to the 7-Tissue Models.

| **Tissue** | **Conductivity (S/m)** |
| --- | --- |
| Skin* | 0.1989 |
| Bone* | 0.0177 |
| Eyes | 1.2000 |
| Cerebrospinal Fluid | 1.6540 |
| Gray Matter | 0.2750 |
| White Matter | 0.1260 |
| Ventricles | 1.6540 |

**Table 2:**
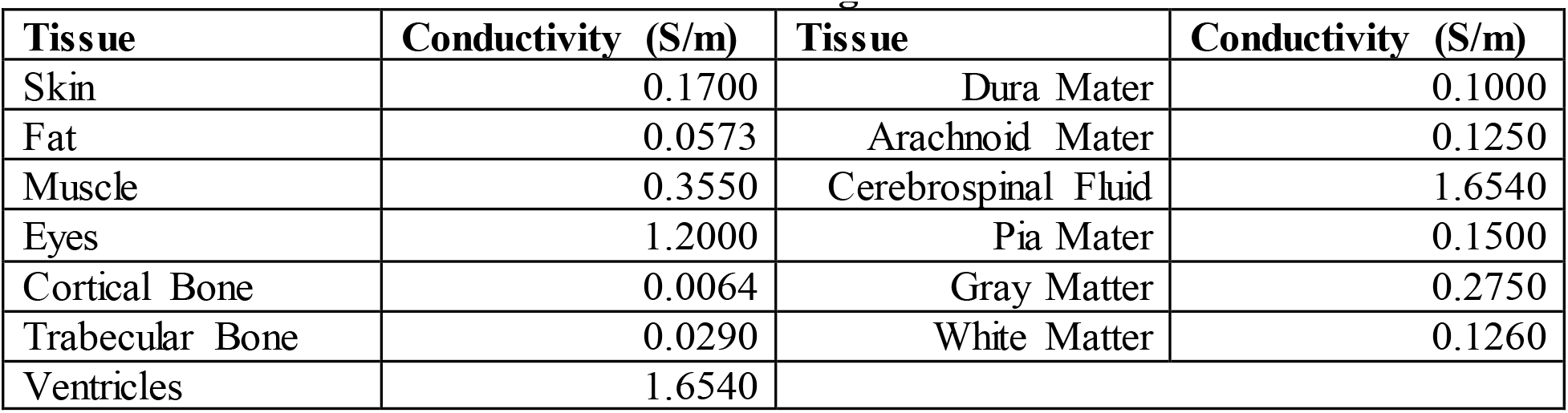
Tissue Conductivities Assigned to the 14-Tissue Models.

### 2.6 Testing Impact of AMR: General Setup

To explore the impact and importance of AMR itself, three distinct quasi-static modalities of forward problem were investigated: TES, TMS, and EEG. Simulations were carried out both on the simple 7-tissue models and on the complex 14-tissue models. In all cases, the source either targets (TES, TMS) or originates in (EEG) the left motor hand area (M1_HAND_).

For each model and each forward problem mode, three solutions were computed. The initial solution was found by solving the model as-is, without invoking AMR. The second solution was found after subjecting the model to AMR with the following parameters: refinement rate = 1% of facets per step, maximum number of refinement steps = 30, and E-field (TES, TMS) or voltage (EEG) in the observation region changes by less than 1% from one AMR pass to the next. The final solution was computed after subjecting the adaptively-refined model to a final global refinement step, where every facet in the adaptively-refined model was indiscriminately subdivided into four sub-facets. These solutions will be referred to respectively as the “standard” (STD), “adaptive mesh refinement” (AMR), and “reference” (REF) cases.

Errors were computed between the STD/REF, STD/AMR, and AMR/REF solutions in manners appropriate for the mode of forward problem. The first error describes the amount of improvement possible due to adaptive mesh refinement. The second describes the improvement achieved by applying adaptive mesh refinement with the stated configuration. The third describes the remaining available improvement that could be achieved through a higher refinement rate or stricter convergence criterion (greater number of AMR steps).

Most simulation parameters were consistent across the problem classes, and they were set to extremely conservative values to minimize sources of error unrelated to the adaptive mesh refinement method. Table 3 summarizes these common simulation parameters, together with typical values that may be used for simple or difficult problems as a reference.

**Table 3:** Simulation parameters common to all forward problem classes.

| Parameter | Selected value | Typical value,<br>simple problem | Typical value,<br>difficult problem |
| --- | --- | --- | --- |
| FMM precision | 1e-6 | 1e-2 | 1e-4 |
| GMRES target residual, initial | 1e-5 | 1e-4 | 1e-3 |
| GMRES target residual, general | 1e-5 | 1e-4 | 1e-3 |
| GMRES max iterations, initial | 100 | 20 | 50 |
| GMRES max iterations, general | 50 | 20 | 50 |

The FMM precision was set to 1e-6, where a value of 1e-2 is sufficient for typical problems and 1e-4 is usually used for difficult problems. For almost all invocations of GMRES, the termination criteria were (a) relative residual of 1e-5 or (b) 50 iterations elapsed (a condition which was required when dealing with the 14-tissue models). Since GMRES is invoked on every adaptive step and the solution vectors are rolled forward from step to step, the GMRES convergence usually saturates over the course of the adaptive mesh refinement method, provided that the refinement method itself is converging. To support a fair comparison with the refined solutions, it is required that GMRES convergence must also saturate for the initial non-adaptive solution. For this reason, the maximum number of GMRES iterations for the initial (NA) solution was set to 100.

Further information on the mode-specific stimulus and evaluated error metrics is given in the subsequent sections.

### 2.7 Testing Impact of AMR: Transcranial Electrical Stimulation

To model transcranial electrical stimulation, five voltage electrodes were placed on the skin surface in a focal ring configuration [31] above the motor hand area. The electrodes were circular with radii of 5 mm, and the four return electrodes were separated from the central active electrode by 30 mm center-to-center. Fig. 3a shows the problem geometry for Connectome Subject 122620.

**Fig. 3:**
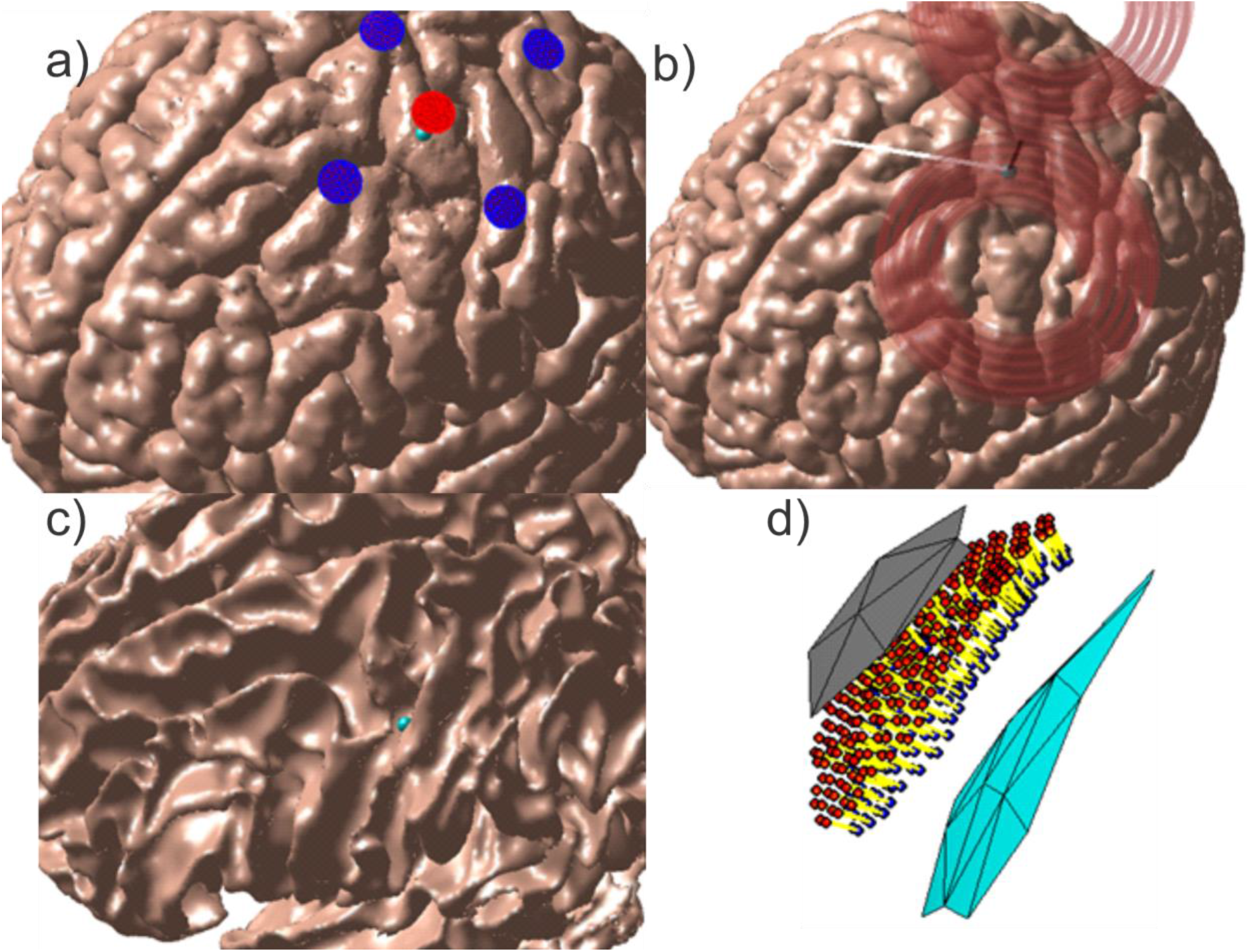
Source configurations for Connectome Subject 122620 (segmented by the headreco pipeline) for each class of forward problem. (a): TES electrode configuration. A focal ring configuration with one positive (red) and four negative (blue) electrodes is shown positioned above the motor hand area. The cyan sphere denotes the selected target point on the GM surface. (b): TMS coil position above the motor hand area at the gray matter surface. The coil model in use is a MagVenture C-B60. The black line denotes the centerline of the coil, the cyan sphere denotes the selected target point on the GM surface, and the white line denotes the expected primary E-field direction at the target point. (c, d): EEG cortical current dipole configuration. (c): Center of cortical dipole cluster (cyan sphere) shown above the WM surface. (d): 260 finite-length dipoles are placed between the GM (gray) and WM (cyan) surfaces centered on the target point. Current flows along the dipoles from the red endpoints to the blue endpoints along the yellow segments.

For the initial solution and all subsequent adaptively-refined solutions, the charge solution was first found for an applied potential of +1 V on the central active electrode and -1 V on the four return electrodes, then linearly scaled to achieve an injected current of 1 mA on the central electrode. Injected current was computed according to

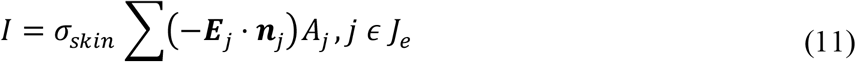

where σ_*skin*_ denotes the conductivity of skin, *J*_*e*_ denotes the set containing the indices of all facets belonging to the central electrode, ***E***_*j*_ denotes the electric field just inside the skin surface at model facet *j*, ***n***_*j*_ denotes the unit normal vector of facet *j* pointing out of the skin surface, and *A*_*j*_ denotes the area of facet *j*. To rescale the charge solution and achieve an injected current of 1 mA, the computed charge density vector **c** was multiplied by 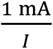. This method grants control over the total injected current without enforcing a spatially-constant current density over the electrode surface, since the spatially-constant field may be unrealistic except for purpose-built electrodes.

A local preemptive refinement step of two iterations of 4:1 subdivision was carried out on skin and electrode facets within a sphere centered on the central electrode and large enough to enclose all electrodes. The skin surface (including electrodes) was then excluded from further adaptive or global refinement. This preemptive refinement was performed because the voltage electrode formulation requires application of a dense preconditioner to couple the Neumann and Dirichlet components of the integral equation, and unsupervised adaptive mesh refinement of the voltage electrodes can quickly grow this preconditioner to an unwieldy size. A useful side effect for the purpose of this study is that the preemptive refinement step prevents modification of the immediate source of injected current by the AMR method; this improves parity with the TMS results (whose coils’ current filaments are never adaptively subdivided) and the EEG results (for which the cortical dipoles are never rearranged, subdivided, or otherwise altered in density).

For the purpose of evaluating the AMR’s E-field convergence criterion, an observation region in the vicinity of the GM target point was constructed. Within a radius of 2 cm from the target point, observation points were placed halfway between the GM and WM surfaces (i.e. on the midlayer) with density approximately equal to the triangular mesh nodal density. The total electric fields from all three solutions were computed at these observation points. Inter-step errors in the E-field were evaluated by applying the L21 norm given in Eq. 12 to these lists of E-field measurements. This error norm, as well as the relative difference measure (RDM) given in Eq. 13, was also applied in the post-simulation analysis of error between the STD/REF, STD/AMR, and AMR/REF solutions.

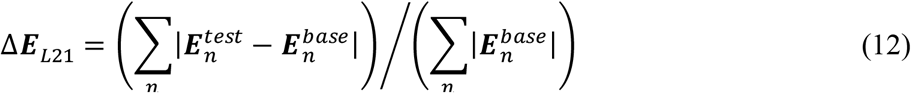

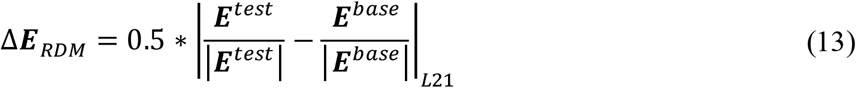

In Eq. 12, |∗| denotes the Euclidean 2-norm, and ***E***_*n*_ denotes the electric field measured at the ROI observation point with index *n*. In Eq. 13, the |∗| operators in the denominators of the test and base terms refer to the matrix 2-norm, and the |∗|_*L*21_ operator indicates that the difference of the test and base terms is to be taken in the L21-norm sense of Eq. 12. Broadly speaking, Δ***E***_*L*21_ measures the relative change in overall E-field magnitude, while Δ*E*_*RDM*_ measures a joint change in spatial distribution of E-field strength and E-field direction. All errors were computed using the total electric field.

### 2.8 Testing Impact of AMR: Transcranial Magnetic Stimulation

To model transcranial magnetic stimulation, a model of a C-B60 coil (MagVenture, Denmark) was positioned according to a target placed on the motor hand area at the GM surface. The coil was placed such that its centerline was normal to the skin surface and passed through the target point on the GM surface, the angle between the fissure longitudinalis and the dominant E-field direction along the coil’s centerline was approximately 45 degrees, and the shortest distance from any part of the skin to any part of the coil windings was 10 mm to account for the coil housing. The coil was modeled as a collection of 17k elementary current segments, and the incident electric field was calculated in terms of these elements’ magnetic vector potential. The coil was driven by a sinusoidal current of amplitude 5 kA and frequency 3 kHz, and the problem was solved at the instant when the time derivative of that current was maximized (94 kA/ms). Fig. 3b shows the problem and coil geometry for Connectome Subject 122620.

The ROI for this method was the same as for TES: observation points were placed at the midlayer surface halfway between GM and WM within a sphere of radius 2 cm centered on the GM target point. Convergence was again evaluated using the L21 norm of the total E-field sampled in the ROI.

### 2.9. Testing Impact of AMR: Electroencephalography

The EEG problem is configured differently from the TMS and TES problems in terms of the location of the sources, definition of the observation region, and quantity measured at the observation region. The sources used in this study are finite-length current dipoles (a current source and current sink) placed roughly halfway between the GM and WM surfaces (cortical layer III/IV) within a sphere of radius 2.3 mm centered on a target point (a wall of the central sulcus) at the motor hand area. The dipoles are oriented roughly normal to the cortex and are assigned a current density following the Okada-Murakami constant of 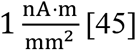 [45]. Fig. 3c shows the dipole location and distribution for Connectome Subject 122620.

The observation region is defined as the set of centroids of all facets belonging to the skin surface, and the field quantity to be evaluated at this surface is the electric potential instead of the E-field. The convergence error metric applied in this case was the 2-norm error given in Eq. 14 below, and the RDM error metric given in Eqs. 15a-b was used for subsequent analysis.

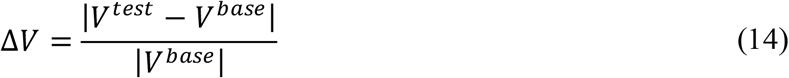

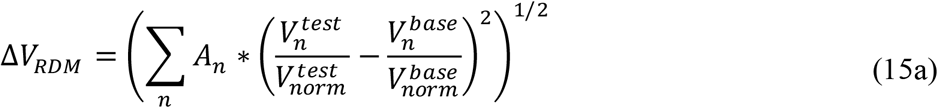

where

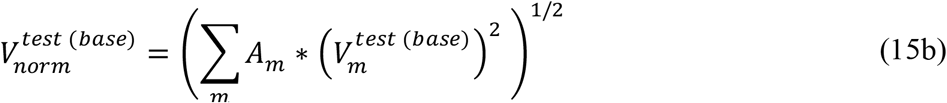

In Eq. 14, *V*^*test*^ and *V*^*base*^ are the vectors of voltages measured at the centroids of all skin surface facets and |∗| denotes the Euclidean vector norm. In Eqs. 15a-b, *A*_*n*_ denotes the area of ROI facet *n*, 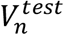 denotes entry *n* of *V*^*test*^, 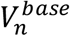 denotes entry *n* of *V*^*base*^, and *n* and *m* iterate over all facets in the ROI.

To achieve good convergence, the 14-tissue EEG test cases required two deviations from the standard treatment applied to the other cases. First, the pia mater surface was removed from the models due to the complicated interaction between the finite-length dipole sources and the double-layer of GM/pia mater charges separated by less than 0.1 mm. Second, the refinement rate was increased from 1% to 3% of model facets per refinement step.

## 3. Results

### 3.1. Impact of AMR: Transcranial Electrical Stimulation

Fig. 4 shows the final refinement maps for the bone, CSF, GM, and WM meshes of Connectome Subject 122620 for the TES test. The color scale denotes the number of subdivisions that were applied in the construction of a given facet in the final model. For example, a facet colored light blue is the product of one 4:1 subdivision step; its edge length is equal to its original (parent) facet’s edge length divided by 2 and its area is equal to its parent’s area divided by 4. An orange facet in this particular figure is the product of three consecutive 4:1 subdivision steps; i.e., its edge length is equal to the original (great-grandparent) facet’s edge length divided by 8, and its area is equal to the original facet’s area divided by 64. Facets in dark blue have not been subdivided at all in the course of the adaptive mesh refinement method.

**Fig. 4:**
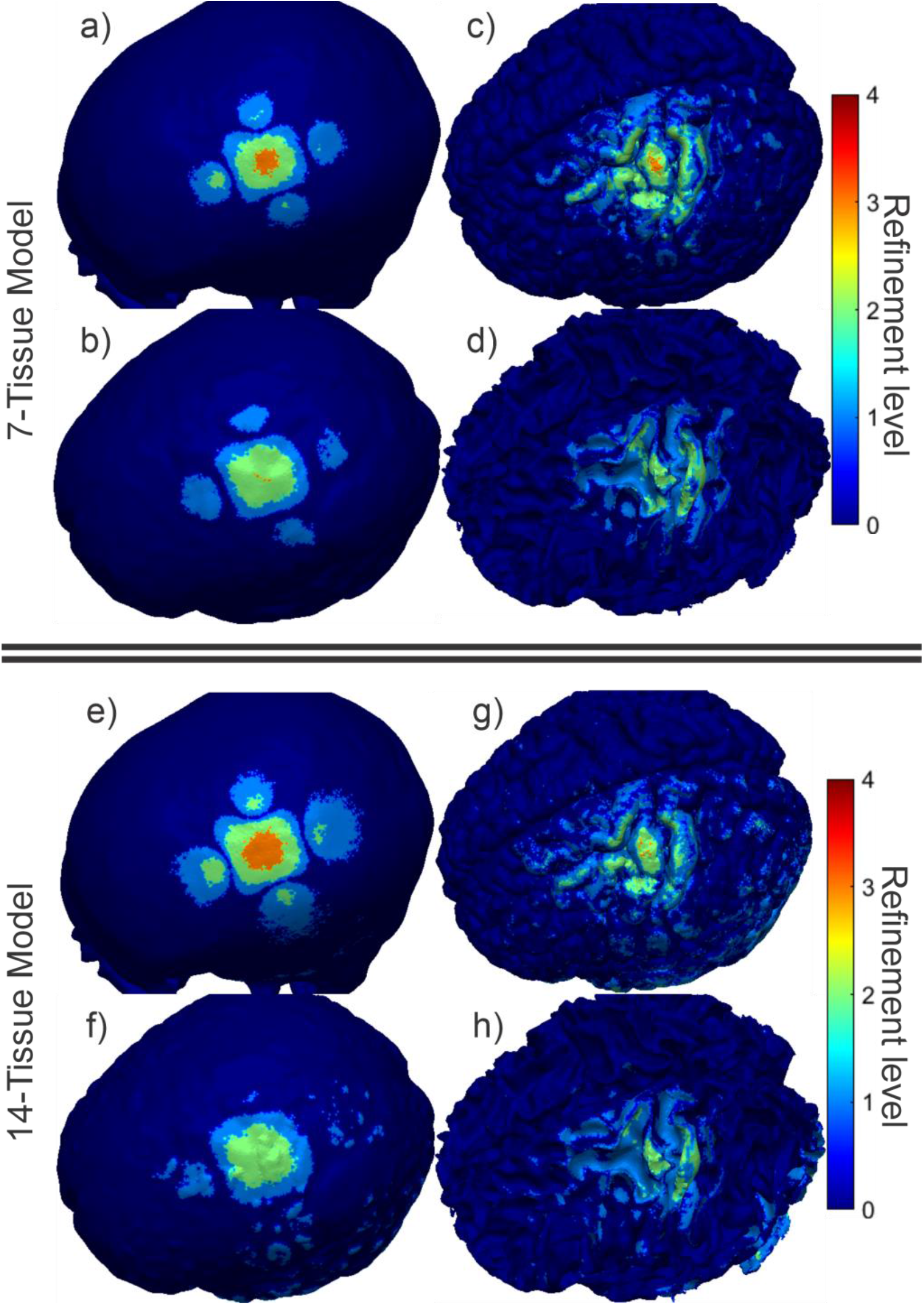
(a-d): TES AMR maps for bone, CSF, GM, and WM (respectively) for the 7-tissue model of Connectome Subject 122620. The color map indicates the number of refinement steps that were applied to subdivide a facet of the initial model into a given facet of the refined model. The current paths beneath the electrodes are clearly visible in the refinement levels of the skull and CSF. (e-h): AMR maps for the same tissues for the 14-tissue model.

Fig. 5 shows the electric field magnitudes in the observation region as well as element -wise absolute differences in the field magnitudes across refinement levels for the 7-tissue and 14-tissue models of Connectome Subject 122620. Fig. 6 presents several summary convergence metrics for all 16 subjects: the number of AMR passes elapsed to achieve convergence by subject, the average (over 16 subjects) and maximum inter-pass charge and E-field errors, and the average and maximum number of GMRES iterations required for each adaptive refinement pass. Finally, Table 4 (Section 3.4) summarizes and compares observation region L21 and RDM errors with the other modalities. Errors were computed on a per-model basis; the table presents the average of those errors over the 16 models in each class. The individual model errors are presented in Appendix A.

**Table 4:** Average (over 16 subjects) summary errors for TES, TMS, and EEG.

| Model and problem | STD/REF<br>(L21, RDM) | STD/AMR<br>(L21, RDM) | AMR/REF<br>(L21, RDM) |
| --- | --- | --- | --- |
| 7-tissue, focal TES | 68.12%, 4.21% | 64.87%, 4.49% | 3.34%, 1.19% |
| 14-tissue, focal TES | 165.54%, 62.92% | 174.20%, 63.37% | 10.37%, 3.58% |
| 7-tissue, TMS | 1.44%, 0.46% | 0.69%, 0.27% | 1.02%, 0.32% |
| 14-tissue, TMS | 3.53%, 1.95% | 1.83%, 0.68% | 2.36%, 1.47% |
| 7-tissue, EEG | 99.67%, 49.37% | 100.13%, 48.62% | 2.67%, 1.94% |
| 13-tissue, EEG | 101.37%, 57.58% | 100.12%, 60.03% | 8.48%, 6.98% |

**Fig. 5:**
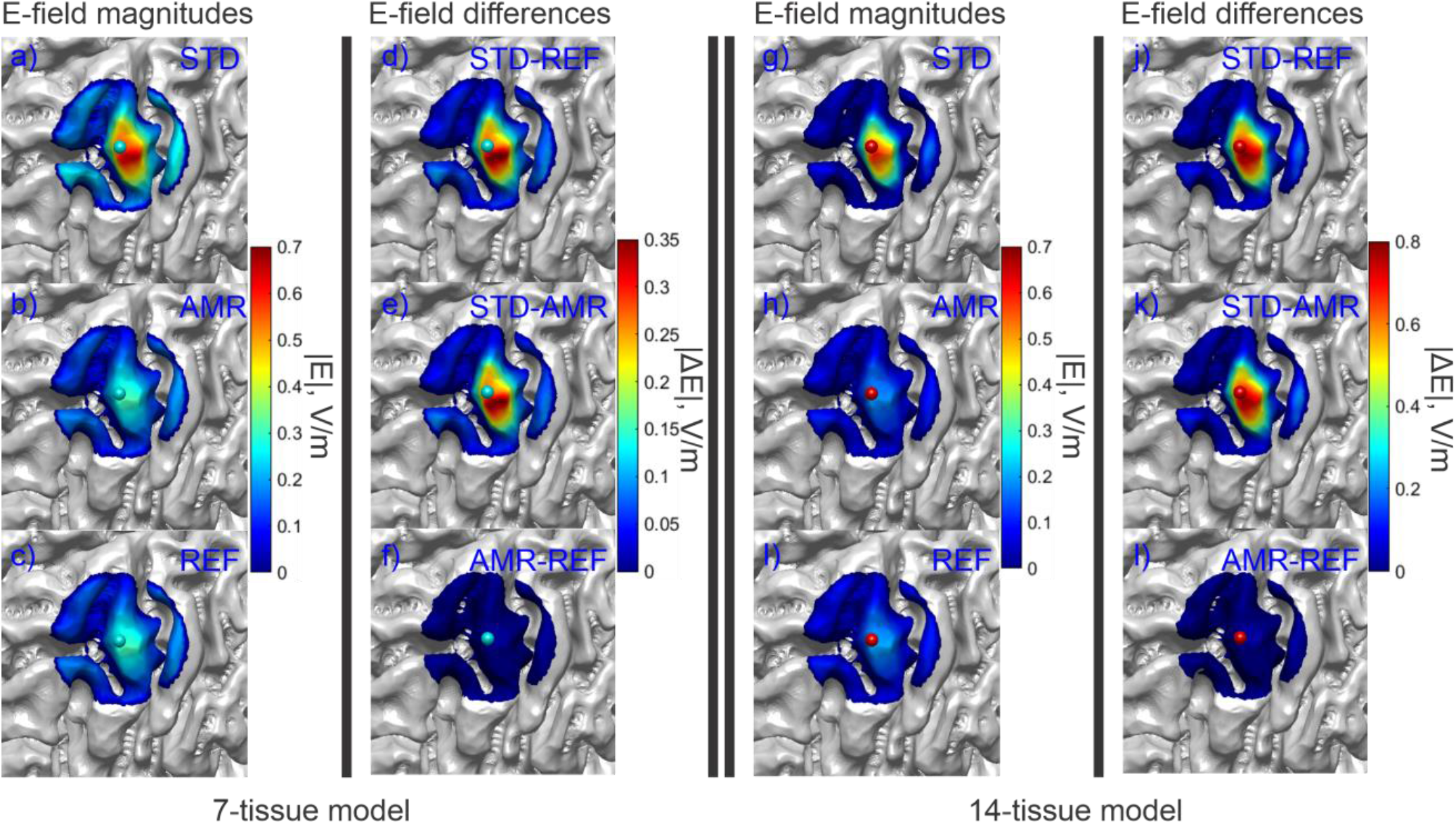
(a-c): Standard, adaptive, and reference (respectively) solutions for the total E-field magnitude (V/m) in the observation region for the 7-tissue model of Connectome Subject 122620. (d-f): Absolute error in E-field is shown between STD/REF, STD/AMR, and AMR/REF solutions respectively. (g-l): Solutions and differences for the 14-tissue model.

**Fig. 6:**
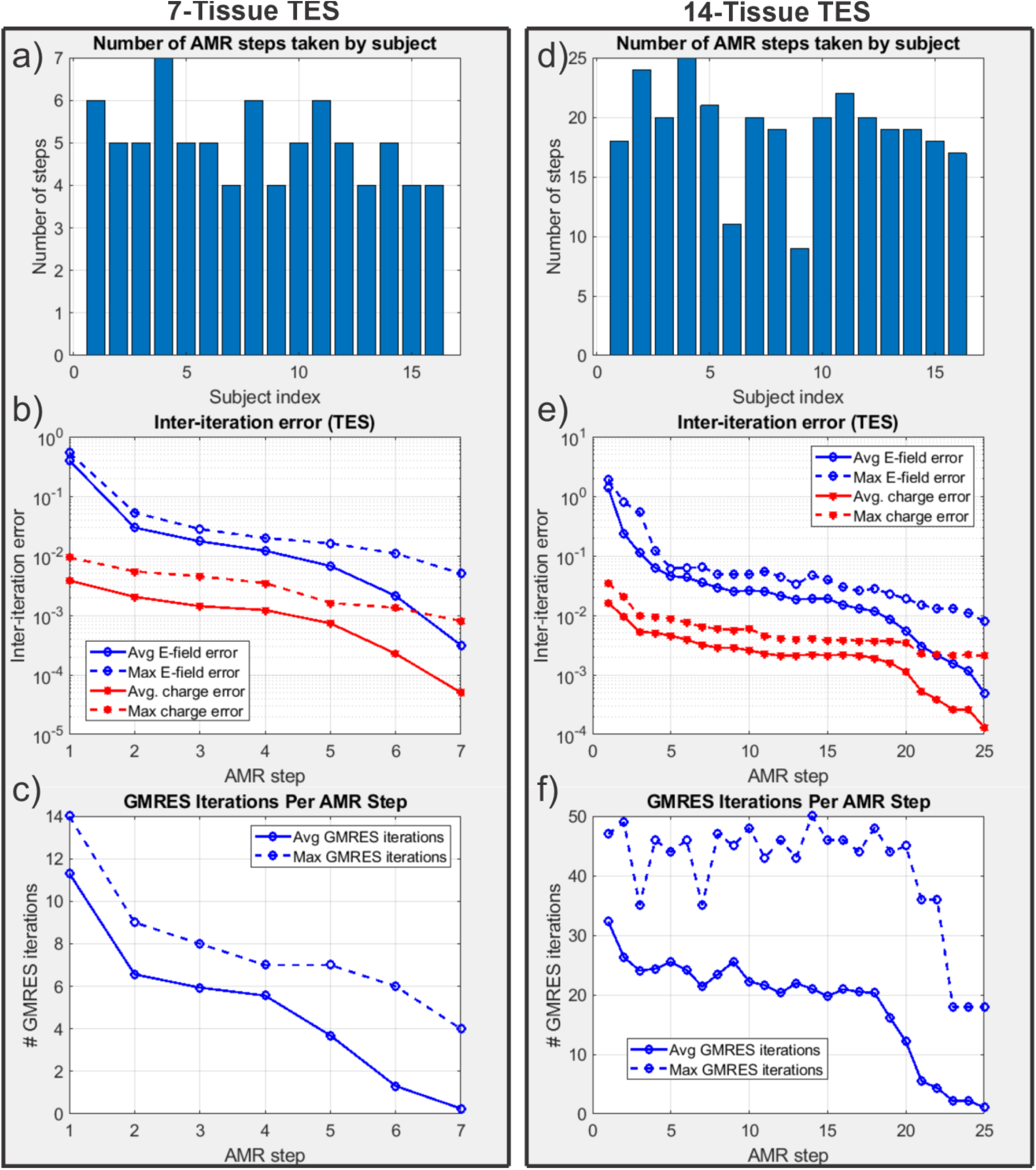
(a): Number of adaptive refinement steps taken by each subject to achieve convergence for the 7-tissue models under TES. (b): Average (16 subjects) and maximum (16 subjects) inter-step E-field and charge solution vector errors at the end of each adaptive mesh refinement step. (c): Average and maximum GMRES iterations required by each AMR step. Note that the average includes entries of 0 for models that had converged prior to the given step; this decision was made to give a reasonable average runtime estimate for large numbers of models. (d-f): The same information is presented for the 14-tissue models under TES.

Note that the number of elapsed AMR passes shown in Fig. 6 differs from the maximum refinement level shown in Fig. 4. The reason for the discrepancy is that only 1% of triangles (per our choice of r = 1%) are subdivided on each step according to the total-charge-based cost function. If a facet were refined 11 consecutive times (i.e., on every AMR pass for the 14-tissue model of Connectome Subject 122620), this would imply that its total charge had been in the top 99^th^ percentile on each of the 10 previous steps, in addition to the start of the 11^th^. By the start of the 11^th^ pass, the facet’s area and corresponding weight would have been reduced by a factor of 4^10^ ≈ 10^6^. Except for facets extremely close (e.g., on the order of nanometers) to sources, such a small facet is not likely to remain in the 99^th^ percentile for total charge on all adaptive passes, and other facets would be selected in its place.

### 3.2. Impact of AMR: Transcranial Magnetic Stimulation

Fig. 7 shows the refinement map for the bone, CSF, GM, and WM meshes of Connectome Subject 122620 for the TMS test. Fig. 8 shows the electric field magnitudes in the observation region as well as element-wise absolute differences in the field magnitudes across refinement levels for the 7-tissue and 14-tissue models of Connectome Subject 122620. Fig. 9 provides convergence summary results for both model classes, Table 4 presents aggregate error metrics over all subjects, and subject-specific errors are presented in Appendix A. It appears that TMS is a rather trivial case where the initial resolution of the model is usually sufficient, even with 14 tissues.

**Fig. 7:**
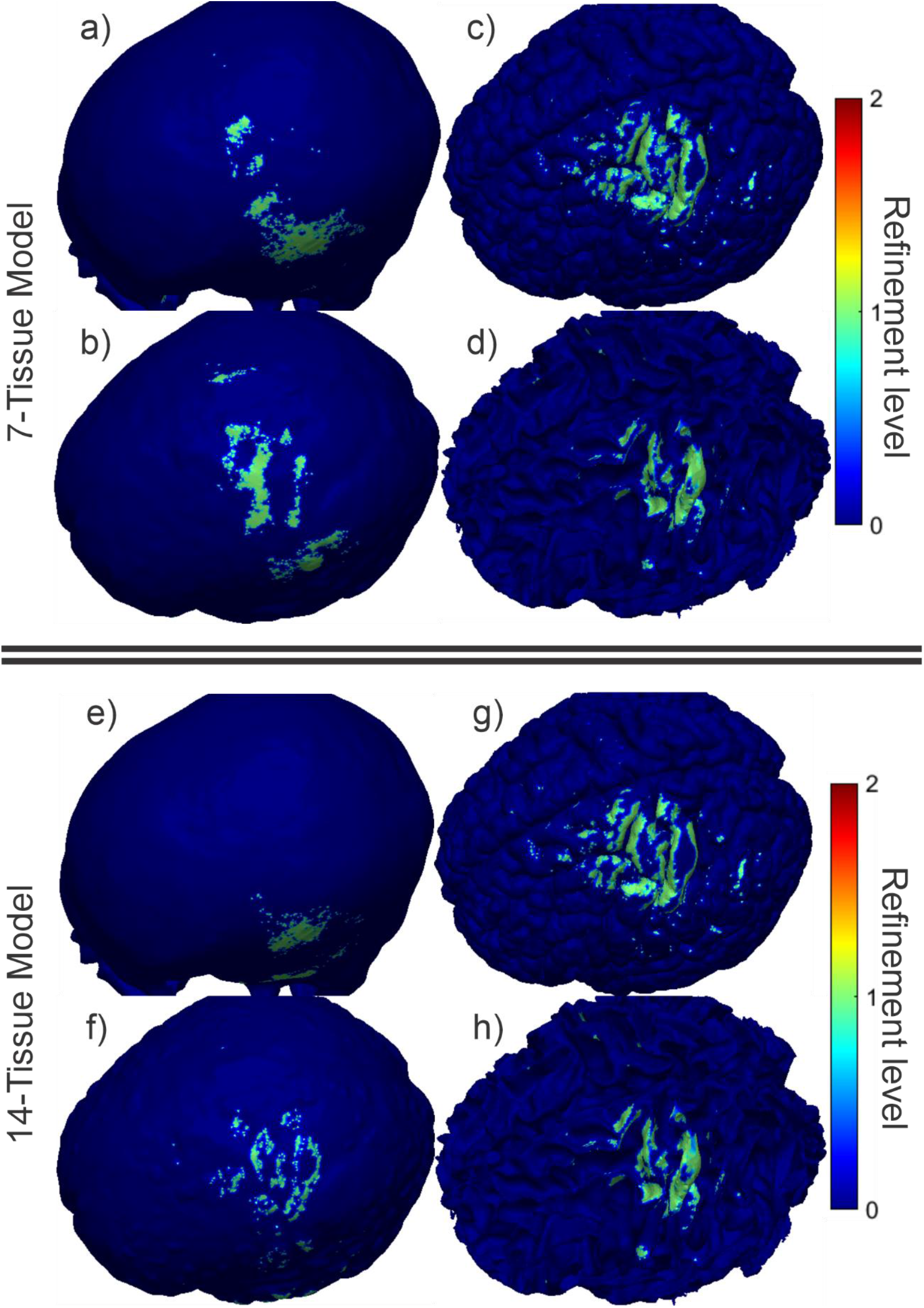
(a-d): TMS AMR maps for bone, CSF, GM, and WM (respectively) for the 7-tissue model of Connectome Subject 122620. The color map indicates the number of refinement steps that were applied to subdivide a facet of the initial model into a given facet of the refined model. Note that the most refinement occurs at the sulcal walls, where the normal component of the total electric field is strongest. (e-h): AMR maps for the same tissues for the 14-tissue model.

**Fig. 8:**
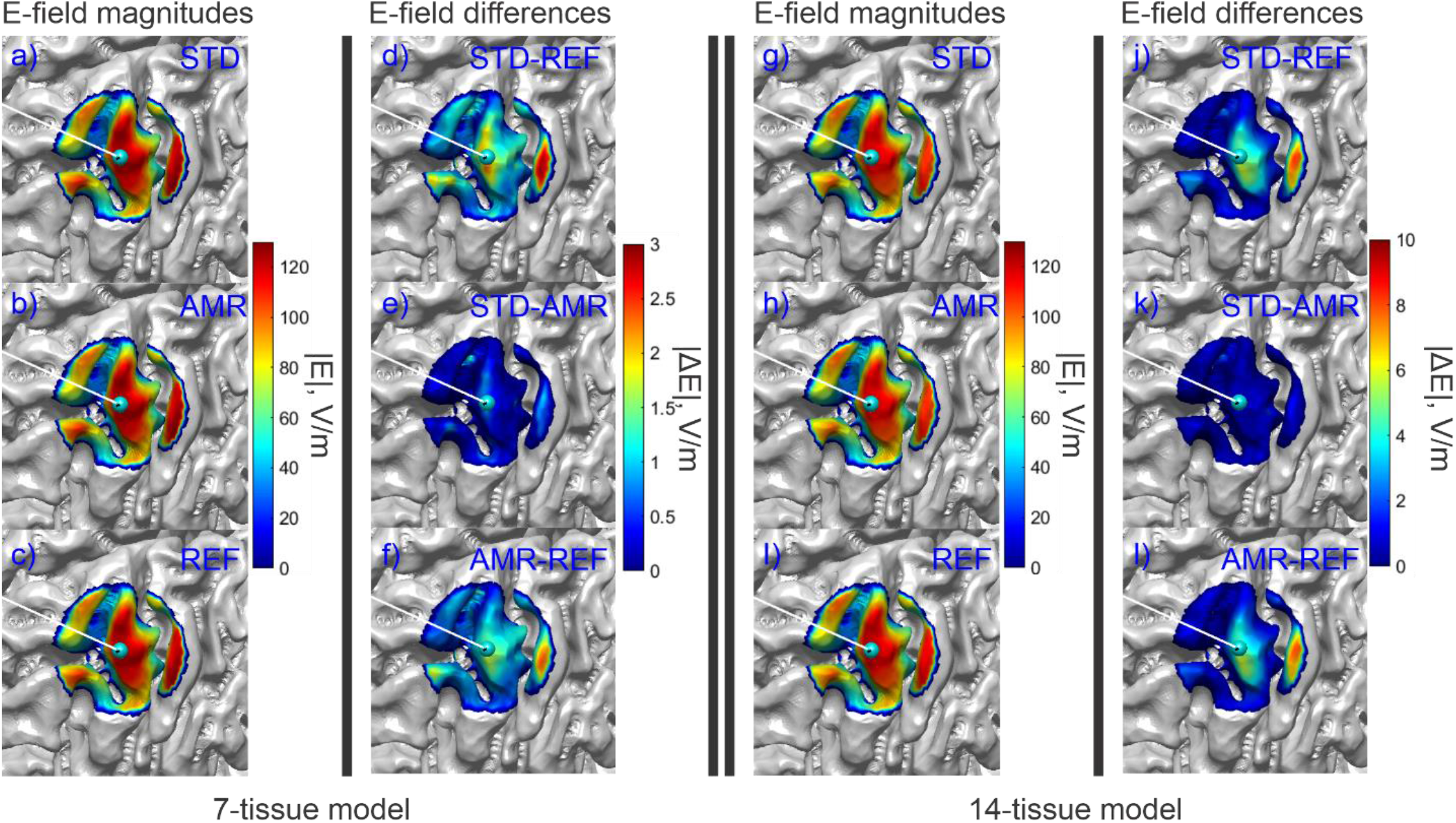
(a-c): Standard, adaptive, and reference (respectively) solutions for the total E-field magnitude (V/m) in the observation region for the 7-tissue model of Connectome Subject 122620. (d-f): Absolute error in E-field is shown between STD/REF, STD/AMR, and AMR/REF solutions respectively. (g-l): Solutions and differences for the 14-tissue model.

**Fig. 9:**
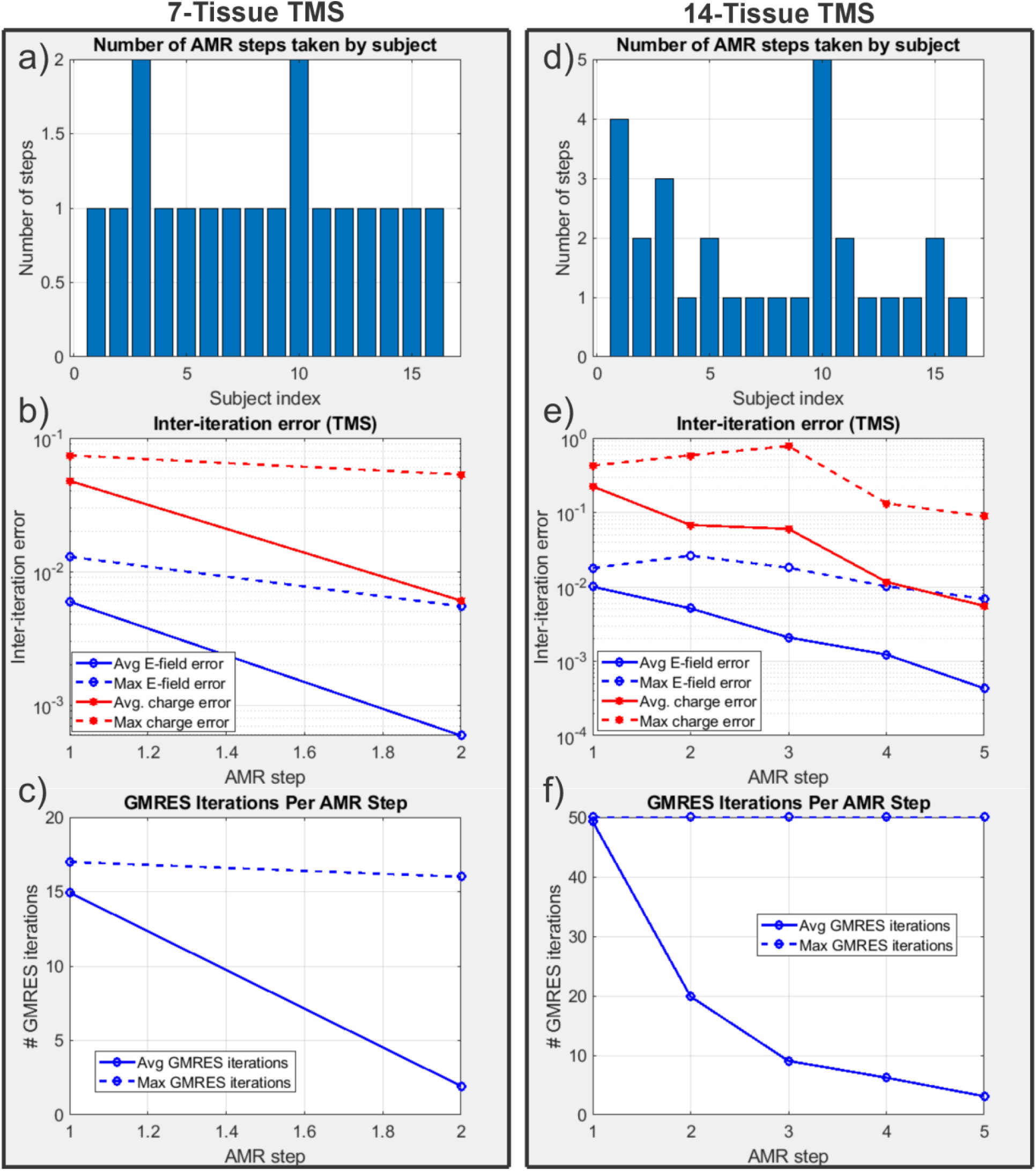
(a): Number of adaptive refinement steps taken by each subject to achieve convergence for the 7-tissue models under TMS. (b): Average (16 subjects) and maximum (16 subjects) inter-step E-field and charge solution vector errors at the end of each adaptive mesh refinement step. (c): Average and maximum GMRES iterations required by each AMR step. (d-f): The same information is presented for the 14-tissue models under TMS.

### 3.3. Impact of AMR: Electroencephalography

Fig. 10 shows the refinement maps for the skin, bone, CSF, GM, and WM meshes of Connectome Subject 122620 for the 7-tissue and 14-tissue EEG tests. Fig. 11 shows a visual comparison of the voltage measured on the skin surface for the same subject as well as element-wise differences in that voltage across the refinement levels. Fig. 12 captures convergence metrics, Table 4 captures aggregate error metrics, and Appendix A captures subject-specific errors.

**Fig. 10:**
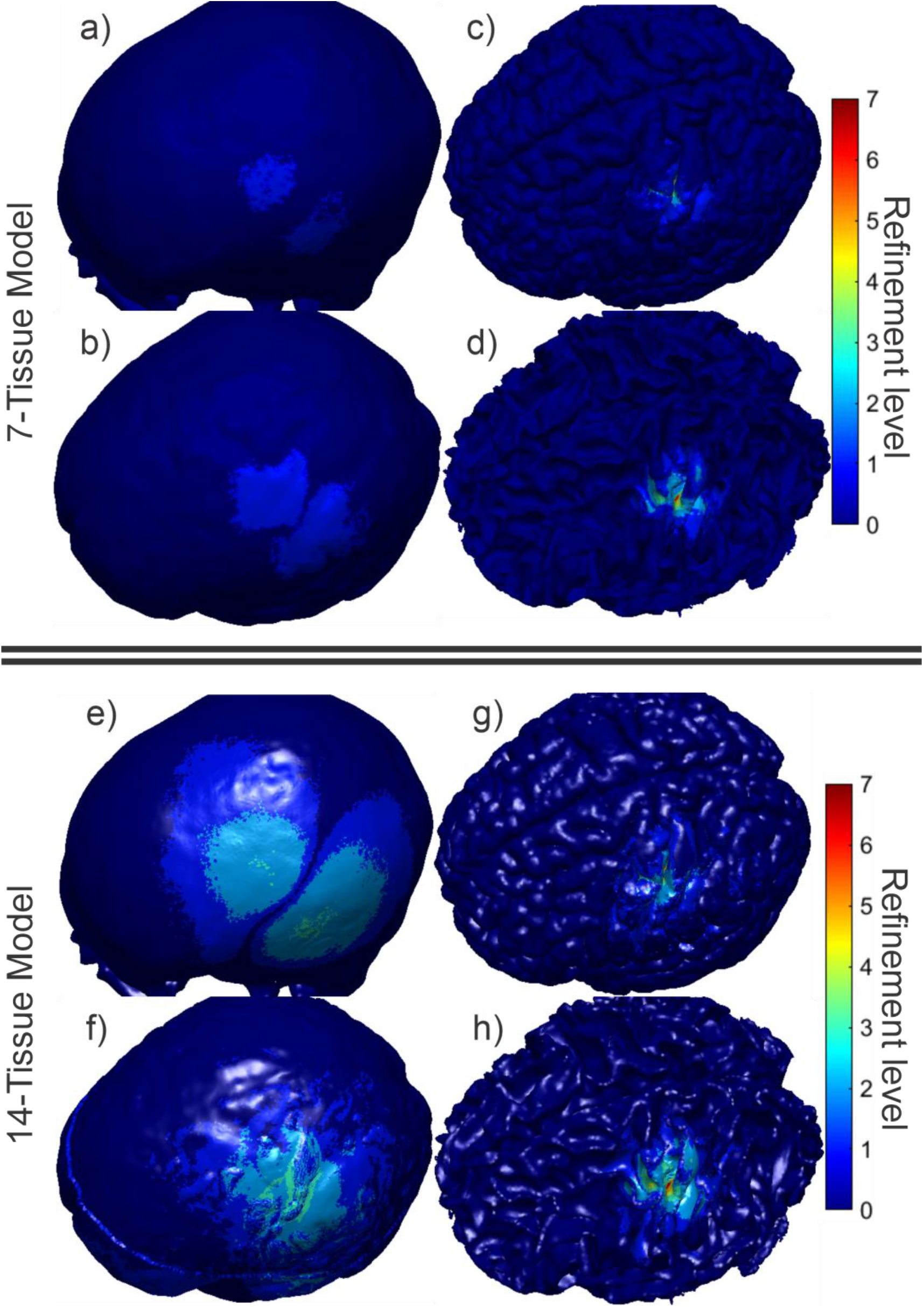
(a-d): EEG AMR maps for bone, CSF, GM, and WM (respectively) for the 7-tissue model of Connectome Subject 122620. The color map indicates the number of refinement steps that were applied to subdivide a facet of the initial model into a given facet of the refined model. The strongest refinement by far occurs in the immediate vicinity of the current dipoles. (e-h): AMR maps for the same tissues for the 14-tissue model.

**Fig. 11:**
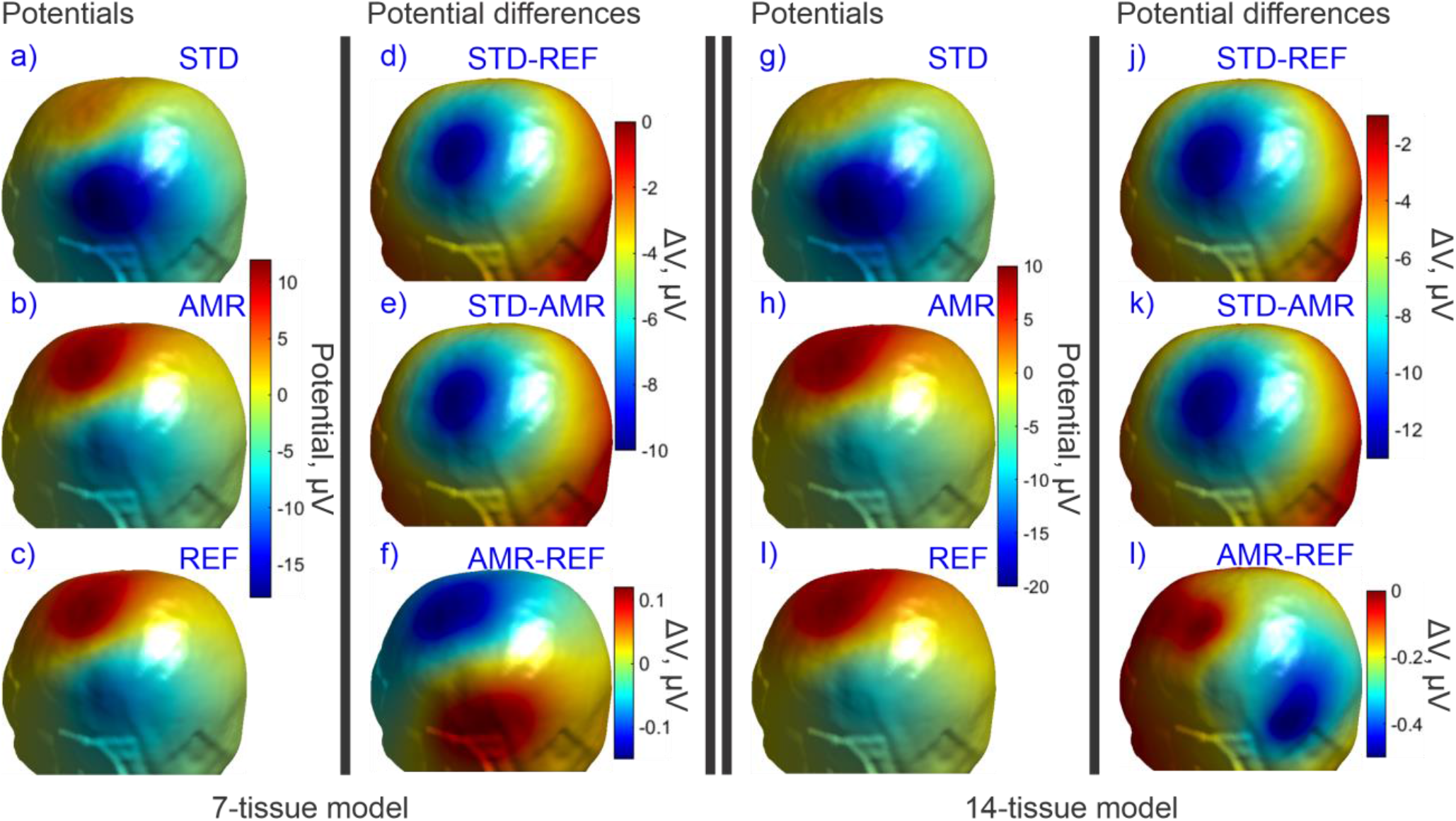
(a-c): Standard, adaptive, and reference (respectively) solutions for the potential (μV) in the observation region for the 7-tissue model of Connectome Subject 122620. (d-f): Signed error in potential is shown between STD/REF, STD/AMR, and AMR/REF solutions respectively. Note the difference in colorbar scales between (d-e) and (f). (g-l): Solutions and differences for the 14-tissue model.

**Fig. 12:**
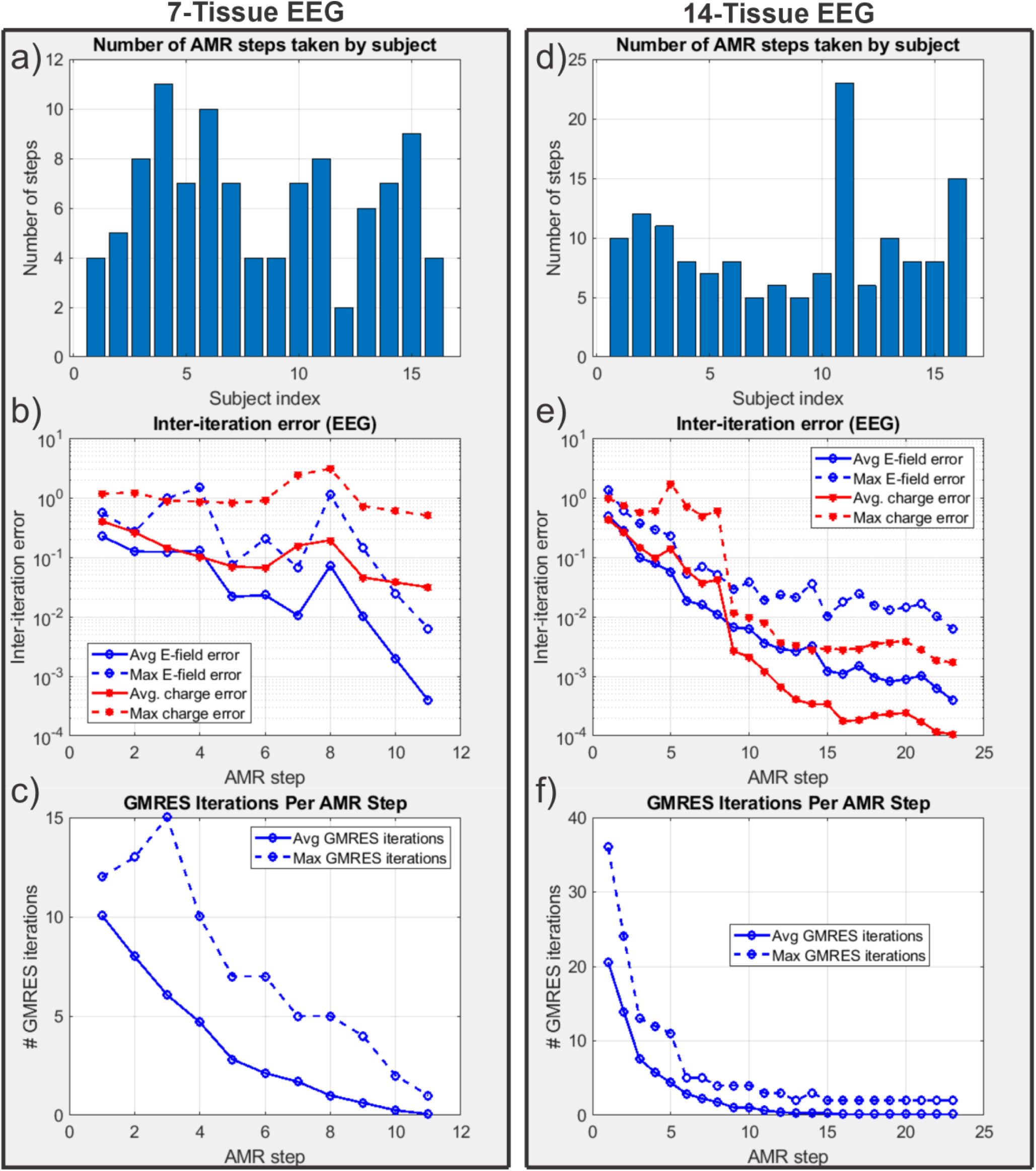
(a): Number of adaptive refinement steps taken by each subject to achieve convergence for the 7-tissue models under EEG. (b): Average (16 subjects) and maximum (16 subjects) inter-step potential and charge solution vector errors at the end of each adaptive mesh refinement step. (c): Average and maximum GMRES iterations required by each AMR step. (d-f): The same information is presented for the 13-tissue models under EEG.

Table 4 summarizes the 2-norm errors computed for the 7-tissue and 14-tissue models for the voltage evaluated over the skin surface. The average error over 14 models is given; per-model errors are presented in Appendix A.

### 3.4: Performance Summary

Table 4 captures multiple summary errors for the three stimulation modes and the two classes of model. As stated previously, the error between the STD and REF cases describes the magnitude of improvement possible due to AMR. The error between the STD and AMR cases describes the improvement that was achieved by AMR, and the error between the AMR and REF cases describes the further improvement that would be possible if the AMR method were carried out for a greater number of iterations or using a higher refinement rate.

Table 5 summarizes the average model size increases that were required to achieve the convergence conditions of the AMR method for each problem class. Recall that the 13-tissue EEG problem was carried out with a refinement rate of 3% of facets per AMR step rather than the 1% rate used in all other methods; the required model size increase for this problem class should make the motivation for that decision apparent. Additionally, the average and standard deviation of number of AMR steps (over 16 subjects per problem class) are reported.

**Table 5:** Average (over 16 subjects) model changes for TES, TMS, and EEG.

| Model and problem | Avg. model size<br>increase | Avg. AMR steps | Std. dev. AMR steps |
| --- | --- | --- | --- |
| 7-tissue, focal TES | 12.64% | 5.0 | 0.9 |
| 14-tissue, focal TES | 67.70% | 18.9 | 4.1 |
| 7-tissue, TMS | 2.87% | 1.1 | 0.3 |
| 14-tissue, TMS | 5.17% | 1.8 | 1.2 |
| 7-tissue, EEG | 21.12% | 6.4 | 2.5 |
| 13-tissue, EEG | 128.16% | 15.8 | 9.9 |

## 4. Discussion

For TMS simulations, there appears to be little benefit from using adaptive mesh refinement to model the electric field arising at the midlayer surface. The main reason for this is that the primary electric field from the coil typically dominates the secondary field by a factor of 2 to 1 or greater. No matter how well the mesh is refined, the coil’s incident field directly dictates the charges induced at the interfaces as well as two thirds of the total electric field at the observation surface. By reciprocity, it is also expected that MEG modeling would not benefit substantially from adaptive mesh refinement in this format since the magnetic field of the localized current sources is unaffected by the conductivity interfaces. Even a full-model pass of 4:1 barycentric subdivision that increases the model size by 300% only results in roughly a 4% E-field magnitude deviation for the 14-tissue model.

In stark contrast, for TES simulations, adaptive mesh refinement becomes very important for accurate E-field simulations. In this case, the primary electric field is zero (except for electrode facets in the case of constant-current electrodes), and the secondary electric field both arises from and dictates the final distribution of the interfacial charges. The electric field at the observation surface has two orders of dependency on the underlying charge distribution: first, the charge-to-charge interaction must properly distribute the charges, and second, the charges must have sufficient resolution to accurately recover the secondary electric field at the observation surface. High mesh resolution is required along the main current paths to accurately model the current distribution. For example, the total current flowing into the GM of Connectome Subject 122620 decreased by 21% over the course of the AMR for the focal TES electrode montage. An appropriately high-resolution mesh is necessary to prevent even minute current deviations due to discretization error. This is highlighted by the results shown in Tables 4 and 5: a 13% (on average) increase in number of unknowns, appropriately distributed by the AMR method, results in a correction to E-field magnitude at the midlayer surface on the order of (on average) 65%. For the 14-tissue TES case, the E-field direction also was subject to a change in direction and/or relative distribution of magnitude of 64%, though a model size increase of 68% was necessary to achieve sufficient resolution in this case.

EEG forward problems also benefit substantially from adaptive mesh refinement, for similar reasons to the TES case. In the case of EEG, the sources are highly singular dipoles that lie within millimeters of model boundaries. Similarly to TES, most current sourced by the cortical dipoles will shunt back to their negative ends without crossing the GM/CSF boundary, let alone reaching the skin. Discretization error in the vicinity of the dipoles can dramatically alter current paths in this critical region and produce radically different voltage distributions at the skin surface . This case is illustrated in Fig. 11. The 7 and 13-tissue models each demonstrate large changes in skin surface voltage magnitude and distribution as captured by Tables 4 and 5. Similarly to TES, the 7-tissue EEG case achieves such dramatic changes after only a 21% increase in number of unknowns on average.

The mesh refinement algorithm as implemented has one very significant limitation: the mesh refinement step does not improve the fidelity of the mesh to the underlying geometry. As previously stated, all subtriangles introduced by the adaptive mesh refinement method are coplanar to their respective parent triangles. As a result, the method strictly introduces additional unknowns into the geometry specified by the initial mesh. Further, charges tend to accumulate at sharp edges in the mesh. When every triangle’s normal vector points in a unique direction with respect to its neighbors, this phenomenon’s impact is minimized. When large regions of locally-coplanar triangles border other large regions of locally-coplanar triangles, charges may accumulate disproportionately along the lines of triangles attached to the border. This accumulation represents a deviation from the underlying problem geometry.

Apart from the obvious desire to use a more accurate model, several approaches may be adopted to minimize these effects. One option is to interpolate the local mesh curvature and translate new vertices to lie upon the interpolated surface. Another is to apply smoothing methods after interpolating additional vertices, although this option risks altering the geometry of the initial mesh. A memory-intensive solution may be to develop a very high-resolution reference mesh and resample new vertices from that reference mesh during refinement. This particular solution would have the added benefit of enabling the initial mesh to start with an even lower resolution, saving computational time.

Multiple improvements could be made to improve the convergence and execution time of the proposed AMR method. In many situations, especially the highly singular EEG problems, it is possible for a facet’s total charge after subdivision (1/4 of the total pre-subdivision charge) to remain greater than the threshold selected for subdivision. A natural improvement would be to preemptively apply a second (third, fourth, …) round of barycentric subdivision to such facets, as the subdivision time is negligible compared to the time that must be spent recomputing neighbor integrals and iteratively solving the refined model. Another possible speed improvement could be achieved by only recomputing neighbor integrals for subdivided facets rather than recomputing all integrals. Other metrics for facet selection, such as current flux through faces, may also speed up convergence for certain problem classes.

An open problem is the appropriate selection of the refinement rate *r*. Large values of *r* tend to increase the number of unnecessary unknowns introduced into the problem at each AMR step. When *r* is too small, neighboring facets in critical regions may be refined on (e.g.) alternating AMR steps, causing erratic convergence behavior that can cause the method to terminate early. This is the phenomenon that prompted the increase from *r* = 1% to *r* = 3% for the 13-tissue EEG models.

## 5. CONCLUSION

In this work, we have described and implemented a conceptually simple, yet effective and computationally efficient adaptive mesh refinement method for the quasi-static charge-based boundary element method with fast multipole acceleration (BEM-FMM). We have demonstrated large improvements to the accuracy of electric potential and electric field measurements at observation surfaces for TES/EEG primary field quantities and no degradation of accuracy for TMS/MEG primary field quantities. For standard 7-tissue TES and EEG forward problems, an increase of only 25% in number of unknowns, allocated efficiently by AMR, reveals changes of 65% or more in the electric field or potential at observation surfaces. The present adaptive mesh refinement method is tailored to the BEM-FMM with 0^th^ order basis functions: it takes advantage of the BEM-FMM’s robustness against manifoldness defects to avoid a full remeshing procedure, thus saving time and minimizing computational complexity. To our knowledge, other methods do not support this simplification yet.

## 6. ACKNOWLEDGMENTS

Research reported in this publication was supported by the National Institute of Mental Health of the National Institutes of Health under Award Number R01MH130490. The content is solely the responsibility of the authors and does not necessarily represent the official views of the National Institutes of Health.

