## Appendix A for "An Adaptive H-Refinement Method for the Boundary Element Fast Multipole Method for Quasi-static Electromagnetic Modeling"

### Appendix A: Full error reports for all models

The error metrics presented in Section 3 of the main text were averaged over all 16 models. This appendix presents the set of error metrics for each of the 16 models individually as a series of bar charts.

##### A.1. Full error report: Transcranial Electrical Stimulation

Fig. A1 below compares the errors computed for the 16 7-tissue and 14-tissue models in terms of L21 and RDM errors for the total electric field in the observation region under the TES problem class.

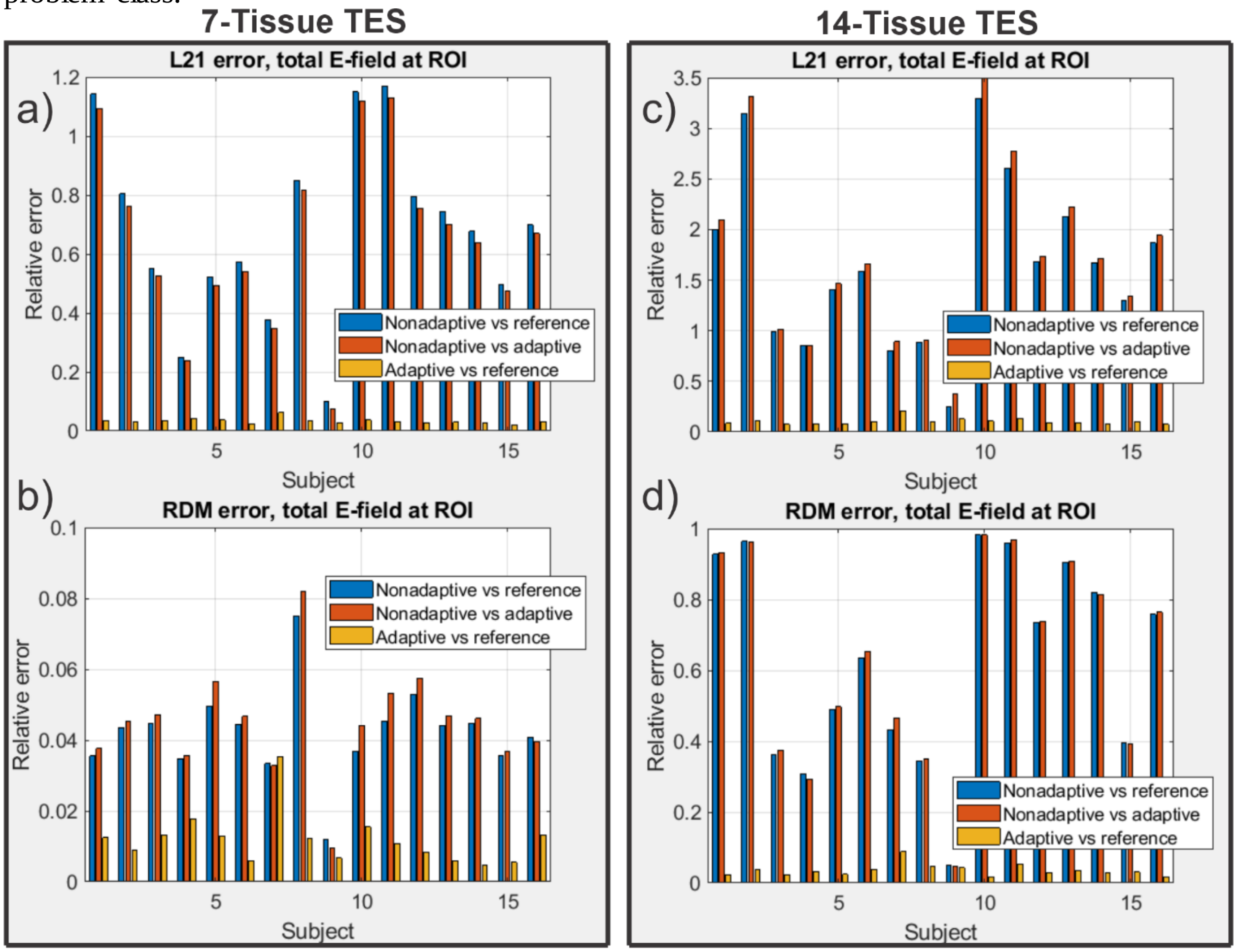


*Fig. A1: L21 (a, c) and RDM (b, d) errors for the 7-tissue (a, b) and 14-tissue (c, d) models under the focal TES forward problem. Blue bars denote the comparison between the nonadaptive solution versus the reference solution (amount of error that could be eliminated by AMR). Red bars denote the comparison between the nonadaptive solution versus the adaptive solution (amount of error that was eliminated by AMR). Gold bars denote the comparison between the adaptive solution and the reference solution (amount of error remaining to be eliminated by, e.g., switching to a stricter convergence criterion or a greater refinement rate). Note the differing relative error scales. Subject 122620, referenced in the main text, is Subject 6 on these graphs.*

**A.2. Impact of AMR: Transcranial Magnetic Stimulation**

Fig. A2 below compares the errors computed for the 16 7-tissue and 14-tissue models in terms of L21 and RDM errors for the total electric field in the observation region under the TMS problem class.

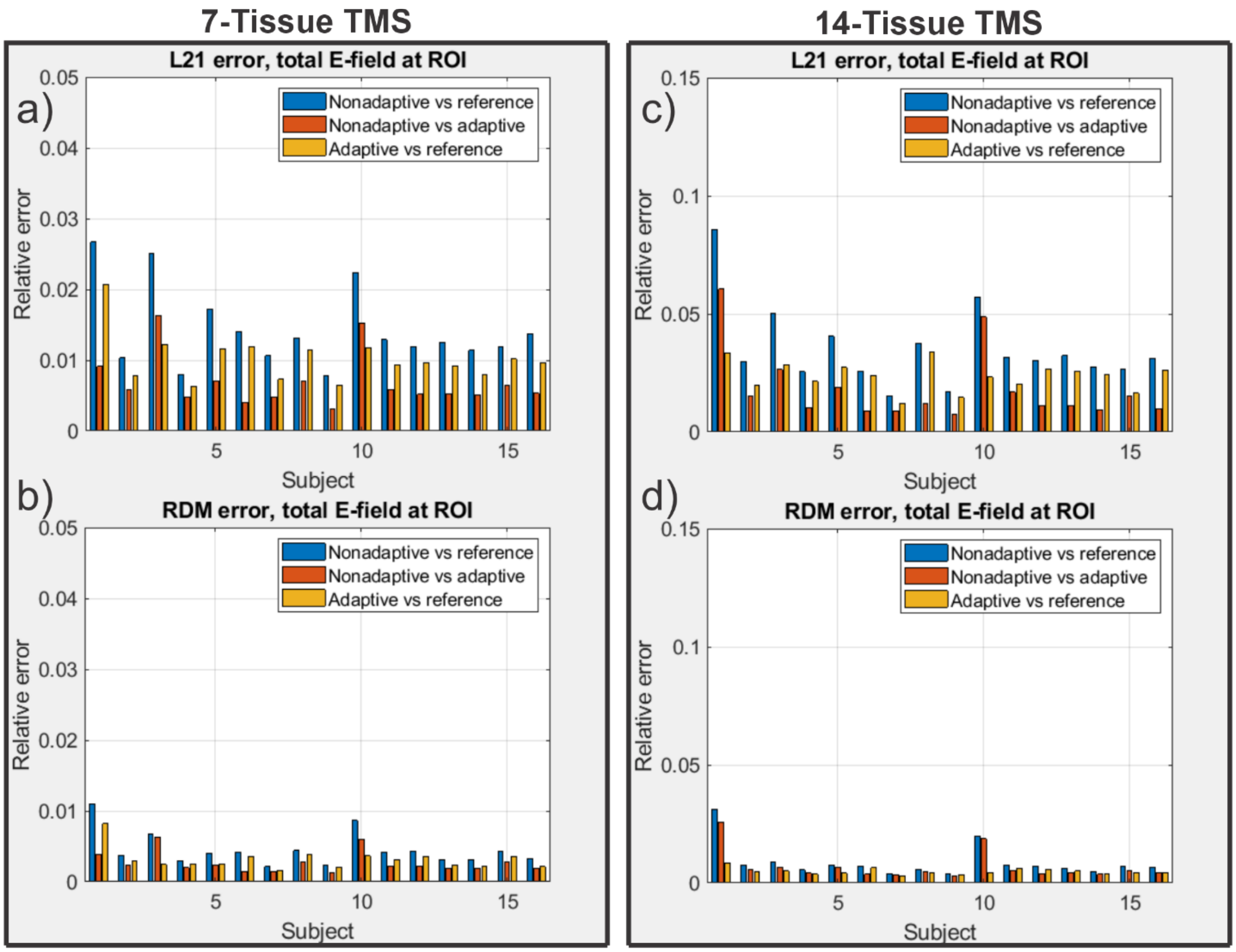


*Fig. A2: L21 (a, c) and RDM (b, d) errors for the 7-tissue (a, b) and 14-tissue (c, d) models under the TMS forward problem. Blue bars denote the comparison between the nonadaptive solution versus the reference solution (amount of error that could be eliminated by AMR). Red bars denote the comparison between the nonadaptive solution versus the adaptive solution (amount of error that was eliminated by AMR). Gold bars denote the comparison between the adaptive solution and the reference solution (amount of error remaining to be eliminated by, e.g., switching to a stricter convergence criterion or a greater refinement rate). Note the differing relative error scales. Subject 122620, referenced in the main text, is Subject 6 on these graphs.*

**A.3. Impact of AMR: Electroencephalography**

Fig. A3 below compares the errors computed for the 16 7-tissue and 14-tissue models in terms of L21 and RDM errors for the potential over the entire skin surface under the EEG problem class.

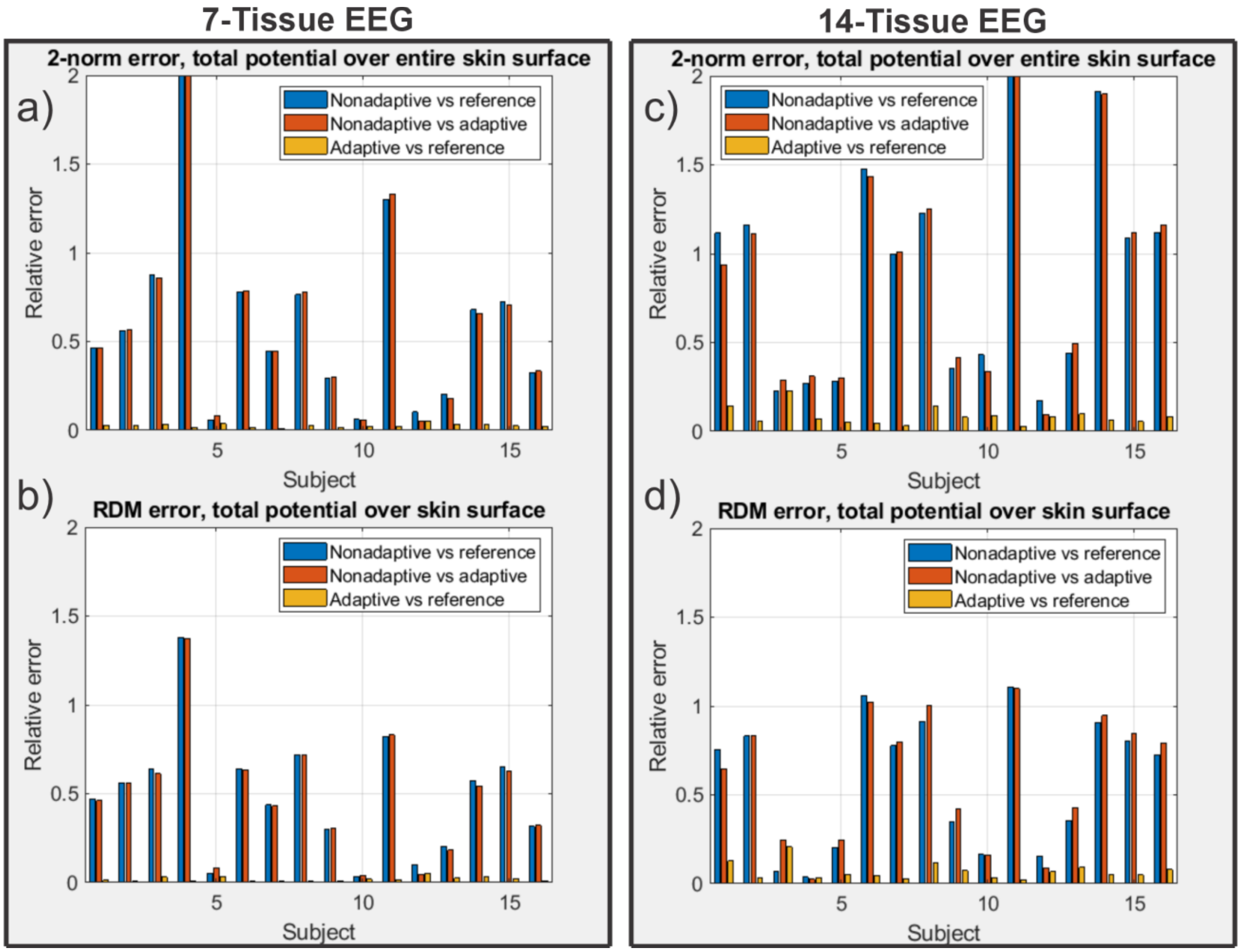


*Fig. A3: 2-norm (a, c) and RDM (b, d) errors for the 7-tissue (a, b) and 14-tissue (c, d) models under the EEG forward problem. Blue bars denote the comparison between the nonadaptive solution versus the reference solution (amount of error that could be eliminated by AMR). Red bars denote the comparison between the nonadaptive solution versus the adaptive solution (amount of error that was eliminated by AMR). Gold bars denote the comparison between the adaptive solution and the reference solution (amount of error remaining to be eliminated by, e.g., switching to a stricter convergence criterion or a greater refinement rate). 122620, referenced in the main text, is Subject 6 on these graphs.*
